# Complex ancient genetic structure and cultural transitions in southern African populations

**DOI:** 10.1101/043562

**Authors:** Francesco Montinaro, George BJ Busby, Miguel Gonzalez-Santos, Ockie Oosthuitzen, Erika Oosthuizen, Paolo Anagnostou, Giovanni Destro-Bisol, Vincenzo L Pascali, Cristian Capelli

## Abstract

The characterization of the structure of southern Africa populations has been the subject of numerous genetic, medical, linguistic, archaeological and anthropological investigations. Current diversity in the subcontinent is the result of complex episodes of genetic admixture and cultural contact between the early inhabitants and the migrants that have arrived in the region over the last 2,000 years. Here we analyze 1,856 individuals from 91 populations, comprising novel and available genotype data to characterize the genetic ancestry profiles of 631 individuals from 51 southern African populations. Combining local ancestry and allele frequency analyses we identify a tripartite, ancient, Khoesan-related genetic structure, which correlates with geography, but not with linguistic affiliation or subsistence strategy. The fine mapping of these components in southern African populations reveals admixture dynamics and episodes of cultural reversion involving several Khoesan groups and highlights different mixtures of ancestral components in Bantu speakers and Coloured individuals.

## Introduction

Southern Africa is characterised by substantial spatial and diachronic cultural variation. Archaeologically, the prehistory of this part of the African continent has been characterized by extended regional variation in lithic industries at the interface between the Middle and Later Stone Ages [Mitchell, 2002]. The different stone-based technologies dated to between 19 and 12 thousand years ago (Kya) have been interpreted in the context of long term isolation, while similarities observed among various Robberg industry stone bladelets found in southern Africa suggested some long-range contact [Mitchell, 2002]. The more recent appearance of pastoralism and agriculture has further complicated the cultural profile of this region. Human and livestock remains documented the arrival of herders in the region less than 2 Kya and several disciplines have attempted to map the local dispersal of agro-pastoralist Bantu speaking populations during the last few centuries. The arrival of European colonists and the subsequent relocation of groups from Asia have added additional complexity to the history of the region. Extended variation can be also observed from a linguistic point of view. Bantu languages are the most commonly spoken in southern Africa, where they have been subdivided into Western and Southern in relation to their geographical distribution and similarities. Some of the non-Bantu languages spoken in southern Africa are characterised by click-sounds and are often referred to as Khoesan (here intended as a non-genealogical group of click-containing languages spoken by a variety of southern African herders and hunter-gatherers [Guldemann and Fehn, 2014]. These languages are classified into three major families [Blench, 2006, Guldemann and Fehn, 2014]: the Kx’a, the Taa and the Khoe-Kwadi, and are characterized by broad and overlapping geographic distributions. This cultural complexity extends also to the different subsistence economies implemented by groups who reside in this region, which include hunter-gathering, animal husbandry and agriculture, plus various combinations of these strategies [Barnard, 1992, Murdock, 1981].

From a genetic point of view, Africa hosts most of the worldwide genomic variability [Campbell and Tishkoff, 2010], and some of the earliest branching Y chromosome and mitochondrial DNA lineages are located in the Southern part of the continent [Tishkoff et al., 2007, Batini et al., 2011, Rosa and Brehem, 2011, Barbieri et al., 2013b]. Due to their potential significance for the origin of modern humans, groups residing in southern Africa have attracted the attention of both geneticists and the general public [Batini et al., 2011, Schlebusch et al., 2012, Pickrell et al., 2012, Barbieri et al., 2013b, Gurdasani et al., 2015]. Such interest has capitalized on the advent of new tools for genome analysis which have contributed to a better characterization and understanding of the history of southern African populations [Henn et al., 2012, Schlebusch et al., 2012, Pickrell et al., 2012, 2014, Kim et al., 2014]. Model-based analyses have demonstrated that populations located north of the Kalahari desert, such as Jux’Hoan and !Xun, are characterized by a so called *Northern* component, which is substantially different from that characterizing populations located to the south of the Kalahari (referred to as the *Southern* component; [Schlebusch et al., 2012, Pickrell and Pritchard, 2012]). However, in-depth analyses of Khoesan genetics have suggested a greater degree of complexity within Khoesan-speaking populations. For example, Schlebusch et al. [2012] highlighted the genetic peculiarity of Gxui and Gxxana individuals when compared with Northern and Southern Khoesan (here referring to the geographic location of Khoesan speaking groups), while Petersen and collaborators [Petersen et al., 2013] suggested additional structure among Northern Khoesan populations (Jux’Hoan and !Xun). In addition to this early structure, a signal of West Eurasian ancestry, which predates the arrival of Bantu-speaking farmers, has also been detected [Schlebusch et al., 2012, Pickrell et al., 2014].

Despite several investigations conducted in the past few years, we are still far from a detailed dissection of the genomic structure related to Khoesan speaking populations. Its exhaustive characterization is challenging due to the fact that various ancestral groups have overlapped over the last millennia and that gene-flow has probably been common among groups. In this context, the legacy left by Khoesan populations in highly admixed groups such as southern African Bantu speakers and Coloured populations is far from clear, which makes the design and the interpretation of regional genome-wide association studies challenging [Rosenberg et al., 2010, Price et al., 2010]. The reconstruction of the ancestry profiles of these populations is further complicated by the fact that groups speaking different languages and implementing different lifestyles have been in contact for extended periods of time, prompting genetic and cultural exchange. The ability to rapidly switch to different cultural packages is potentially an effective solution for surviving in challenging environments [Kinahan, 2001].

Here, to further dissect and clarify the genomic stratification of southern African populations, we analyzed 1,856 individuals from 91 populations using a combination of novel (59 individuals from 7 populations) and published genome-wide SNP data. By applying a local ancestry deconvolution approach we highlight previously unobserved complexity in the Khoesan-related ancestry components and generate novel insight into the genetic history of the region. We provide evidence for the presence of at least three distinct Khoesan ancestral components and reveal a substantial degree of admixture between Khoesan groups. Our fine dissection of the Khoesan-related legacy in highly admixed populations also reveals slight, but significant genetic structure between Coloured and Bantu-speaking populations, suggesting different admixture histories in these two groups [Tishkoff et al., 2009, Schlebusch et al., 2012, Barbieri et al., 2013b, Pickrell and Pritchard, 2012, Petersen et al., 2013, González-Santos et al., 2015, Marks et al., 2015, Busby et al., 2016]. Finally, we demonstrate that Khoesan-related ancestry structure is highly correlated with the geographic location of populations but not with linguistic affiliations or subsistence strategies.

## Results

### Complex population structure and mixed ancestry in southern Africa

We described population structure in southern Africa using the ”Unphased Dataset” (See Methods) comprising 1,856 individuals from 91 populations genotyped at 63,767 autosomal markers (Fig.1, Fig. S1). After kinship analysis, 25 individuals were removed due to the existence of 25 pairs of highly related individuals. Ten of these pairs contained individuals genotyped in different studies: 2 Bantu South Africa from the HGDP [Li et al., 2008] with a high kinship index with 2 Herero from Schlebusch et al. [2012], 3 and 5 pairs among the Jux’Hoan (from Pickrell and Pritchard [2012], Petersen et al. [2013] and Schlebusch et al. [2012]) and the Khomani (from Henn et al. [2011] and Schlebusch et al. [2012]), respectively.

**Figure 1:**
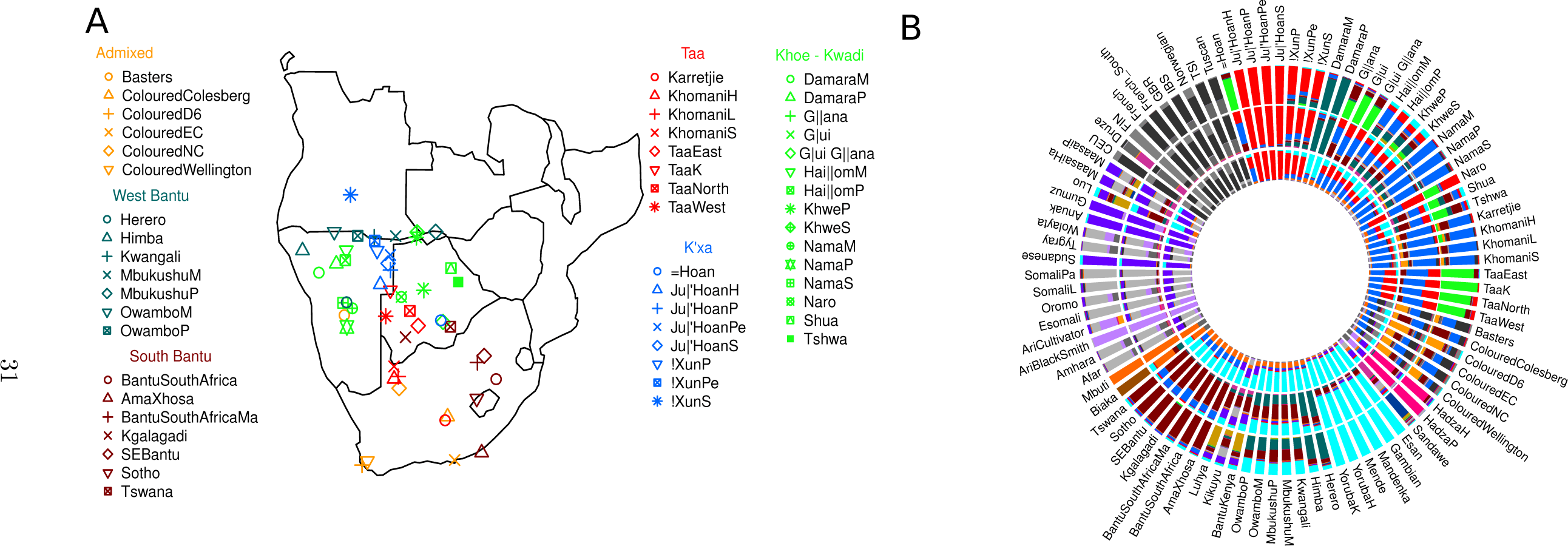
The genetic structure of southern Africa populations. **A.** southern Africa populations analyzed in this study. Different colors are associated with different language/ethnic affiliation. The complete dataset used for analysis is shown in Fig. SI, and Table SI. **B**. Admixture results for *K* = 10, 15, 20 (from the inner to the outer circle). Colors at the center reflects the affiliation shown at Fig. 1A and Fig. SI. We analyzed 1856 individuals for 91 populations and averaged the results in a population based barplot. The full set of results (*K* = {1‥20}) for individuals and populations are reported in Fig. S2 and Fig. S3.

To visualize the evolutionary relationships among the analyzed individuals, we used ADMIXTURE [Alexander et al., 2009], varying the prescribed number of clusters, *K*, from 2 to 20 (Fig. 1, Fig. S2, Fig. S3). At *K* = 7, all African populations are mostly characterized as a mixture of four African-specific components defined by language, geography or ethnicity, representing Khoesan (red and blue), Niger-Congo (turquoise), East African (purple), and rainforest Hunter-Gatherer (Pygmies, orange) populations. Interestingly, the latter is also present in Western and Eastern Bantu populations, and in the Hadza, Sandawe and Maasai from East Africa, possibly reflecting admixture and/or the existence of a geographically extended Pygmy-related ancestral component [Destro-Bisol et al., 2004, Patin et al., 2014].

**Figure 2:**
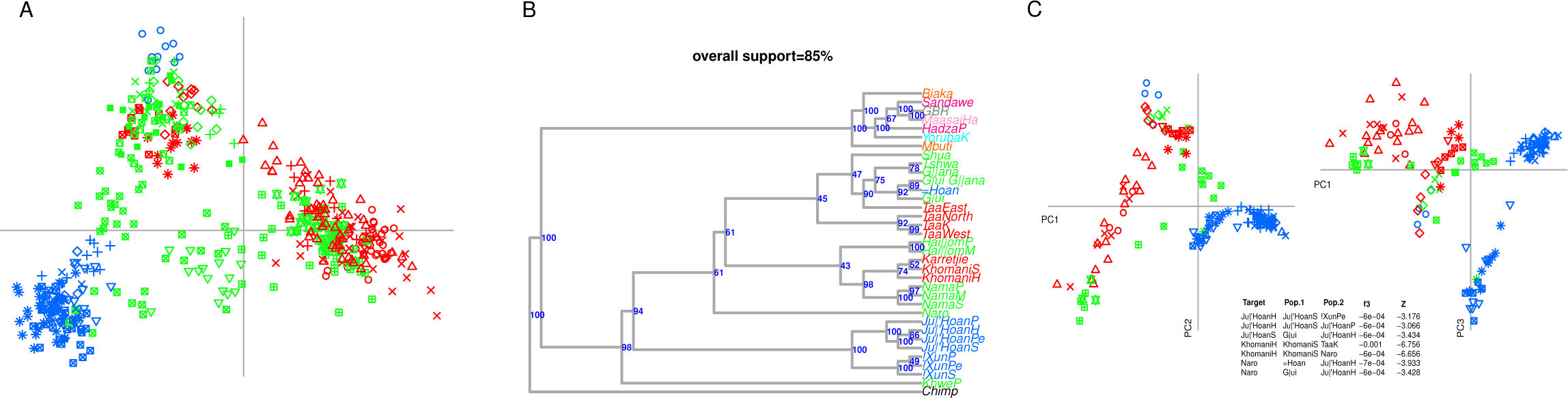
Local Ancestry deconvolution reveals complex Khoesan-related structure. **A.** Principal Component Analyses of Khoesan specific fragments. We extracted fragments with high (>99%) probability to be derived from Khoesan populations and visualized it in a Multi-Dimensional Scaling Plot, as described in Methods section. **B.** Maximum likelihood tree of Khoesan populations. We selected all the Khoesan populations and added 7 African and European populations. We performed ten different run and assessed the support of each tree trough 100 bootstraps (Fig. S6). Colour keys are as in Fig.lA and Fig SI. C. Principal component analysis of individuals with more than 80% of Khoesan-related genetic ancestry. We used the K=3 ADMIXTURE run to select individuals characterised by at least 80% of Khoesan genetic ancestry and performed a Principal component Analysis as described in Methods section. The two most significant *fy* between “Target” and sources (“PI” and “P2”) populations, including standard deviation (sd) and Z-score, are reported.

Among Khoesan groups, signatures of admixture and possible cultural transition are evident in most of the populations. For example the Damara show a high fraction of Bantu-like ancestry components (Fig. 1, Fig. S2, Fig. S3). More generally, almost all of the Khoesan populations show a non-negligible fraction of ancestry components which are modal in East Africa and Europe, consistent with ancient and recent migrations from these regions [Tishkoff et al., 2009, Schlebusch et al., 2012, Pickrell et al., 2014]. At *K* = 8 the Khoesan ancestry component splits into two. One is modal in all the K’xa speaking populations (less in the =Hoan), while the other is common in all the other Khoesan populations, reaching the highest frequency in the Nama and Khomani. At *K* = 9 and *K* = 10 additional Eurasian components emerge, which differentiate from Afro-Asiatic ancestries.

We noted that although the smallest estimated cross-validation (CV) values are found for *K* = 9 and 10 (Fig. S4), analyses performed at higher values of *K* provide insights into the genomic history and substructure of populations, so we describe these results below. At *K* = 12, the component common in Niger-Congo speaking populations splits into two, one common in Western and Central African populations and in Western Bantu speakers, the other more common in Southeastern Bantu speakers, consistent with these representing the last stage of the Bantu migration process [González-Santos et al., 2015]. Interestingly, this Southeastern Bantu component is present in most Khoesan populations from Botswana, Lesotho and South Africa, providing evidence for admixture during the expansion of Bantu-speaking populations [Tishkoff et al., 2009, Schlebusch et al., 2012, Barbieri et al., 2013b, Pickrell and Pritchard, 2012, Petersen et al., 2013, González-Santos et al., 2015, Marks et al., 2015, Busby et al., 2016]. The presence of ancestry related to Western Bantu speakers in some of the Western Khoesan populations such as the Khoe, the Haixxom and the Nama, is consistent with their current geographic position and could be interpreted as a signature of admixture events. At *K* = 13, an ancestry component present only in the recently admixed population of South Africa and Namibia (Coloured and Basters) emerges: *F*_*ST*_ values of this component suggests genetic similarity with European populations. However, the genetic distance between this ancestry component and the other African components is consistently smaller than the values estimated when using European populations (Fig. 5) which suggests that the admixture between African and Eurasian populations might have generated a novel combination of allele frequencies which is now captured by this component (Fig.1, Fig. S2, Fig. S3). We tested this hypothesis by exploring admixture runs performed on different admixed populations composed by a variable fraction of British (GBR) and Yoruba (YRI) ancestry. Interestingly at *K* = 3, in all the simulations the ADMIXTURE software models a third component which is characterized by a mixture of the other two (Fig. S6). At *K* = 14, a component which almost exclusively characterizes Western Bantu populations becomes evident. Interestingly, this component is modal in the Damara, Herero and Himba populations, providing some evidence for a closer affinity [Barbieri et al., 2013a]. In addition, this same ancestral component is at high frequency in all the other Bantu populations, where it is complemented by the presence of the Southeastern Bantu component. At *K* = 15, the Sandawe population differentiates from other groups from Tanzania, Hadza and Maasai.

### Local ancestry analysis reveals three distinct Khoesan-related ancestries

Ancestry analysis of Khoesan populations is complicated by the fact that the genomes of most of the contemporary populations are a mosaic of multiple ancestries [Pickrell et al., 2014, Marks et al., 2015, Busby et al., 2016]. For this reason we performed a Local Ancestry Multi-Dimensional-Scaling (LAMDS) analysis using only genomic fragments assigned with high confidence to Khoesan ancestry (”Khoesan Ancestry dataset”, see Methods). Similar methods have previously been successful in assessing the continental legacy of American populations [Moreno-Estrada et al., 2013]. However, such methods rely on a large set of reference populations that can be used as a scaffold for local PCA visualization, which were not available here. We therefore applied a new LAMDS approach, in which an IBS-based distance matrix is generated by comparing only those variants on chromosomal segments identified as being of Khoesan ancestry. This analysis has the advantage of using chromosomes instead of individuals and allows one to plot admixed populations even when there is not a comprehensive reference dataset. We assessed the possible impact of variable number of markers analyzed through a resampling procedure, as explained in Methods section. The mean correlation of IBS between the whole and the resampled dataset is always higher than 0.8 and reach 0.9 with as little as 1,500 markers, which suggests that the impact of different numbers of markers on IBS estimates is negligible (Fig. S7). In addition, the tract length density for the two dataset is very similar (Fig. S8, with the number of short chunks identified for the Affymetrix dataset being slightly larger than the Illumina. However, given that the missing data are excluded in our pairwise IBS distance estimation, such observation is not expected to cause substantial bias.

Previous analyses of Khoesan populations suggested the existence of two distinct ancestries, although subsequent investigation has pointed to a more complex underlying structure [Schlebusch et al., 2012, Pickrell and Pritchard, 2012, Pickrell et al., 2014]. Our LAMDS analysis reveals three main groups of Khoesan-related ancestry (Fig. 2A). The first group (*Northern* Khoesan) is composed of all the K’xa speaking populations located at the North of the Kalahari(Jux’Hoan and !Xun) with the exception of the more central =Hoan. The remaining two groups are composed by the Nama, Khomani and Karretjie (*Southern* Khoesan) and all of the remaining Khoesan populations (*Central* Khoesan).

We further investigate the presence of the described Khoesan-related three-way genetic structure with a series of analyses of the ”Unphased dataset”. First, we built Maximum Likelihood (ML) trees from allele frequencies using TREEMIX, and tested their robustness with bootstrapping. Khoesan populations (with the exception of the Damara, who were excluded from this analysis due to their low Khoesan ancestry, as reported in Fig. 1B) are separated in three groups, a pattern consistently found across tree bootstraps. The three-way partition described by TREEMIX broadly mirrors the clustering patterns suggested by the ancestry-based analysis (Fig. 2B, S9). The Karretjie seem to be more related to the Khomani than to the Nama, with the Naro acting as an outgroup to these two branches, probably as a result of their complex pattern of Khoesan ancestry. Similarly, Haixxom and Shua form a distinct branch that splits from the other Southern Khoesan. However, the poor support for these branches highlights their complex genetic make-up, which could be the result of interactions and admixture between different Khoesan populations. On the other hand, the split which separates the !Xun samples from the Jux’Hoan is well supported, providing further evidence for genetic distinctiveness between these two K’xa populations [Petersen et al., 2013], an observation which is further suggested by their separation along the third component of the PCA analysis of individuals with more than 80% of Khoesan ancestry (see below; Fig. 2B, C).

PCA using individuals with at least 80% Khoesan ancestry provides additional evidence for the Khoesan ancestry tripartition (Fig. 2C). Specifically, the three vertices of the plot recapitulate the ADMIXTURE and TREEMIX analyses, with the three clusters composed by populations defined by different amounts of *Northern*, *Central* and *Southern* Khoesan ancestries. We note that in the ADMIXTURE analysis described above, at *K* = 16 three Khoesan-related components emerge, separating all the populations from the central Kalahari area (Botswana) speaking Taa, K’xa and Khoe-Kwadi (*Central* Khoesan) from the K’xa in the North (*Northern* Khoesan) and the Nama, Khomani and Karretije in the South (*Southern* Khoesan). Notably, the *F*_*ST*_ values between these three components are similar, suggesting either a deep split (possibly followed by admixture) and/or drastic demographic events, such as bottlenecks or founder effects. Using *F*_*ST*_ and sample sizes for Jux’Hoan and Taa [Kim et al., 2014], we estimate a splitting time of ~25 Kya (95% confidence interval 18-32 Kya among all the *F*_*ST*_ and sample size combinations, Fig. S10A) when ancestral components inferred by admixture are used, which is broadly consistent with previous estimates [Pickrell et al., 2012, Kim et al., 2014]. When pairwise population *F*_*ST*_ values were used (Fig. S10B), and using a generation length of 29 years [Fenner, 2005], the inferred splitting time is 14Kya (2-27Kya), which is probably reflecting the effect of admixture involving Khoesan and/or non-Khoesan populations. Nevertheless, both approaches seem to demonstrate that this genetic structure could have a prehistorical rather than historical origin [Pickrell et al., 2012, Kim et al., 2014]. It is interesting to note that among the Basters and the Coloured, the *Southern* Khoesan component represents most of the Khoesan-like ancestry, whilst, conversely, in South African and Lesotho Bantu-speaking populations the majority component is *Central* Khoesan. Furthermore, a substantial number of Khoesan populations show a combination of these three ancestries, suggesting extensive admixture in the history of these populations. Interestingly, 5 out of the 11 of the *Central* Khoesan populations are closer to *Northern* populations by means of the *f* _4_(*Ju|hoanP e, Nama, X, Chimp*) test, suggesting gene flow between these two groups (Fig. S11). However, the interpretation of these tests are challenged by admixture which skewed allele frequency of the ”source” populations. Alternatively, as suggested by the importance of the geography in the modern day genetic structure of the populations in the area, these results can reflect a genetic ”isolation by distance equilibrium”.

In order to provide a temporal framework on the adxmiture dynamics in the area we used MALDER, which exploit a weight LD decay statistics to infer the time of admixture between populations (Fig. S12). Overall, the results are consistent with similar investigations performed in earlier investigations [Pickrell and Pritchard, 2012, Pickrell et al., 2014, Busby et al., 2016]. Among Khoesan populations, a large number have signatures suggesting two different admixture events; the first, less than 10 generations ago and involving African and non african populations, is concordant with Colonial times in the region, while the second, involving similar sets of populations and dated ~40–60 generations ago (~1160–1740 years ago), is probably related to the arrival of the pastoralists in the area [Pickrell et al., 2012, 2014]. All the Coloured populations share with the Khoesan recent episodes of admixture (~4–7 generations ago, ~116–203 years ago) but this admixture signature has not been found in the Southern-east Bantu, with the exception of BantuSouthAfricaMa. On the other hand, all of them share a admixture event dated ~17–32 gen. ago (~493–923 years ago) which is consistent with the arrival of Bantu speaking populations in the area. The BantuSouthAfricaMa show evidence of 5 admixture events, which could be explained by the fact that the sampled individuals are from different Southern Bantu Speaking populations all sampled in the Soweto-Johannesburg metropolitan area.

We modeled the geography of population structure in Khoesan populations taking advantage of the Bayesian statistical framework implemented in SpaceMix. The resulting geogenetic map which summarizes the genetic structure and the admixture events among populations is shown in Fig. S13A. The results are consistent with the one obtained in the analysis described before, providing further insight about the distribution of the genetic variability in the area. First, the 95% ellipses in the geogenetic map built by the algorithm highlights the existence of three main Khoesan groups, according to their geographic location (Fig.S13A). In addition, this approach is able to detect apparent sub-structure within the three clusters, such as the genetic differentiation between !Xun and Jux’Hoan, or between Khomani and Nama. Interestingly, The *Central* group seems to be further subdivided into a Eastern and Western group, which adds additional evidence for the complexity of the area. Furthermore several admixture events (*a >* 1%) were identified, confirming the existence of past relationships between the three groups. High non Khosean contributions have been inferred for Khomani, !Xun, Jux’Hoan and Naro, among several other minor admixture events involving other populations. The run tracts and the correlation among the inferred and observed parameters suggest that the analysis and the model accurately describes the observed data (Fig. 13B, C, D).

### Contemporary Khoesan populations contain a mixture of Khoesan-related ancestries

Our LAMDS analysis offers further insight into the relationships between Khoesan groups (Fig. 2A). In fact, chromosomes from several populations seem to be scattered between different clusters, potentially as a result of admixture. For example, Khomani individuals are spread towards groups enriched in *Central* Khoesan ancestry, the Naro and some of the Jux’Hoan occupy a position intermediate between populations characterized mostly by *Central* and *Northern* components and the Haixxom are scattered between individuals with *Northern* and *Southern* Khoesan genetic profiles. With the exception of the Haixxom (where all individuals have less than 80% Khoesan ancestry), these results are consistent with our PCA analyses based on the subset of individuals in each group with more than 80% Khoesan ancestry. It is important to note that all the analysis converge towards a tripartite genetic structure in Southern Africa suggest that the errors due to the ”computational phasing” is negligible. In addition it has been previously shown [Hellenthal et al., 2014] that different phasing methods have usually consistent results. Moreover, PCA confirms similar patterns to those described above for the Khomani and Naro, which are spread towards groups rich in *Central* and *Northern* Khoesan ancestry, respectively (Fig. 2C). We formally tested for admixture between populations applying the *f*_3_ analysis on the same dataset (Fig. 2C); among the significant tests, we reported the two most negative Z-score for each population tested; all the comparison are reported in Table S3. Significant *f*_3_ statistics provide evidence that these mixed ancestries are the result of admixture between different ancestral Khoesan populations (Table S3, Fig.2C). None of the *Central* Khoesan populations show significant evidence of admixture between *Northern* and *Southern* groups. In the Khomani, the lowest *f*_3_ values are found when considering Taa populations (Table S3, Fig.2C). The Naro show evidence of admixture involving populations close to Jux’Hoan and a central Khoesan population, such as Taa and Gxui. The Jux’Hoan also show significant *f*_3_ values when tested against !Xun (*Northern* Khoesan) and Naro (*Central* Khoesan). Similarly, the !Xun also show evidence for admixture with the Jux’Hoan.

### Khoesan-related genetic structure in admixed populations

To better visualize the relationships between Khoesan and non-Khoesan populations, including individuals with less than 80% Khoesan ancestry, we plotted all southern African groups with 90% utilization distribution density kernels for the three ancestries, estimated using the KernelUD function in the *adehabitatHR* package [Calenge, 2006]. This approach allows us to explore which of the three ancestry components is present in Bantu-speaking, admixed, and Khoesan groups with high Bantu ancestry, such as the Khwe and the Damara. The Khwe cluster with the sympatric Jux’Hoan and !Xun populations, although some individuals are located closer to populations mostly containing a *Central* Khoesan component, potentially reflecting a non-negligible degree of admixture. The Damara, conversely, seem to be genetically closer to the Khomani and Nama, although they are scattered towards the K’xa populations in the North, in accordance with their geographic location. Interestingly, we identified two Owambo individuals with genomic features related to Jux’Hoan and !Xun populations.

All of the other Bantu-speaking groups – with the exception of the Kwangali, who are closer to Taa and K’xa speaking groups from Botswana (*Central* Khoesan) – are genetically related to the cluster defined by Nama, Karretjie and Khomani (*Southern* Khoesan). We noted that admixed individuals mapping to this cluster appear to highlight a partly structured distribution, since the Bantu populations are located on the upper side of the distribution, while Coloured and Basters are on the lower side (Fig. 3A). To test this hypothesis we used *mclust* to explore the most supported number of clusters, from 1 to 9 inclusive, using either the MDS coordinates or the IBS distance matrix. Using the Expectation-Maximization model based clustering algorithm, we inferred 7 clusters using the MDS coordinates and 9 with the distance matrix and the average probability for each population are shown in Fig. 3B and Fig.S14. In both analyses the populations are mostly defined by the same cluster affiliation, although present in different proportions. An additional minor cluster, related to Bantu-speaking populations from Botswana, present in the Southern Bantu populations, is absent in the Coloured and Basters. Such differences are still evident when the complete distance matrix is considered (Fig.S14). These results are supported by the *f*_4_ test *f*_4_(*Khoesan*1*, Khoesan*2*, X, Chimp*) which shows a marked difference between Bantu and Coloured populations. In details, all the Bantu are not significantly closer to ”Central” (Gxui and Gxxana individuals) or ”Southern” Khoesan (Nama) while Basters and Coloured are closer to Southern Khoesan (Fig.S11C). However, caution must be used in the interpretation of these tests when admixed populations are used as “Sources”.

**Figure 3:**
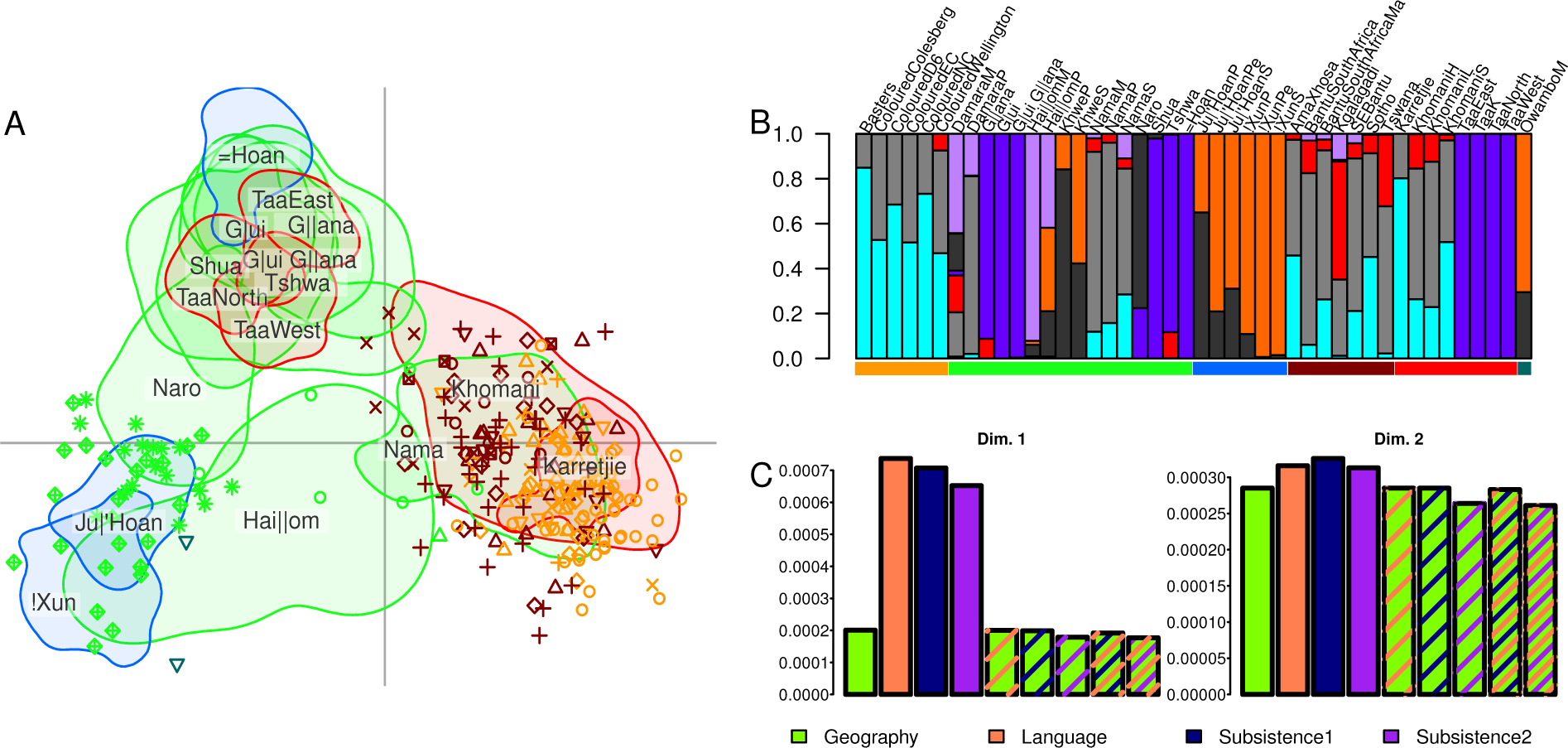
**A.** Genetic structure of admixed southern African populations. In order to provide a simplified version of Figure 2A, we estimated the 90% utilization kernel of Khoesan populations (except Damara and Khwe, see text), and plotted the highly admixed individuals. **B.** Cluster Analysis of genomic fragments. We grouped all the individuals in 7 clusters, as inferred by mclust R package (see methods) and visualized the results in barplot according to populations and language/ethnic affiliation. Colour keys are as in Fig.1A and Fig S1. The results highlight the large heterogeneity in populations sharing the same affiliation and the existence of a slight but significant substructure between Bantu and Coloured populations. **C.** Predictive errors of genetic components for geographic, linguistic and subsistence affiliation or a combination of different covariates (striped bars), for the first two dimensions of the MDS in Fig 2 and 3A (Dim 1 and Dim 2, respectively). Geography better predicts genetic ancestries, though adding new covariates slightly decrease the predictive error.

### Khoesan-related genetic structure and geography, language and subsistence

We performed a Procrustes analysis to test the relationship between genetic and geographic distances and found a statistically significant correlation (Procrustes correlation= 0.65, p *<*0.001), as previously observed by Schlebusch et al. [2012]. We noted that these authors tested this correlation across a small subset of Khoesan populations. Here we extended this analysis to include not only a larger dataset of Khoesan populations but also Khoesan fragments in highly admixed groups such as southern African Bantu-speaking populations and Coloured. To further investigate the association between geography with the observed Khoesan-related structure and to explore the correlation with cultural variables, we evaluated the power of models predicting the positioning of individuals along the two dimensions of the Khoesan-ancestry IBS-based MDS plot (Fig. 3C) for geography, language, and subsistence. Major reductions in model predictive error, which is indicative of better model fit, are only observed when variables are considered in relation to MDS dimension 1 (Fig. 3C), while dimension 2 shows some degree of model prediction reduction only when geography is considered (Fig. 3C). Geography shows the smallest predictive error, and therefore best model-fit, when each variable is singularly considered (geography: 0.000201, language: 0.0007, subsistence1: 0.0007, subsistence2: 0.000652). Although the predictive power of the analysis is improved when multiple variable are considered, the reduction of cross validation error is minimal (Geography + Language: 0.0002, Geography + Subsistence1: 0.000199, Geography + Subsistence2: 0.000179, Geography + Language + Subsistence1: 0.000192, Geography + Language + Subsistence2: 0.000177). These results hint at the existence of a geographically-described ancient complexity in Khoesan ancestry which likely pre-dates the arrival of Bantu and European populations, and which is only marginally captured by current ethno-linguistic population descriptors.

## Discussion

The genetic characterization of populations from the African continent is crucial from an epidemiological, pharmacological, anthropological and evolutionary perspective. Within the continent, southern Africa displays an impressive degree of genetic and cultural diversity, this being a region where groups speak several different languages and implement a variety of different subsistence strategies. From a linguistic point of view, Khoesan languages are unique to this region and are divided in three major families: K’xa, Khoe-Kwadi, and Taa. While the separate grouping of K’xa and Taa speakers has reached a consensus among linguists, the internal structure of the Khoe-Kwadi family is still debated. The most heterogeneous of the three linguistic groups, Khoe-Kwadi, is usually classified into three subgroups; East (spoken by Thswa and Shua), West (Khwe, Gxui, Gllana and Naro) and Khoekhoe, which is currently spoken by the Nama, Damara, and Haixxom populations [Guldemann and Fehn, 2014]. The history of Khoekhoe populations still remains unresolved; for example, the Haixxom and Damara have previously been classified as ’other bushmen’ when their phenotypic, linguistic and/or cultural characteristics were considered [Barnard, 1992]. The Haixxom live in Northern Namibia, and they are thought to be !Xun individuals that have recently acquired the Nama language. The Damara – who were sometimes referred to as BergDama or BergDamara – live in Northern Namibia and their origins are also unclear. Including both herders and foragers, the ancestral population probably arrived in the area before the Nama and Western Bantu populations, such as Herero and Owambo. The arrival of the Nama pastoralists in the Namibia region from an area in the South African Northern Cape (Namaqualand) is a recent event dating to the end of the 19th century [Barnard, 1992]. The first pastoralist populations described by Dutch colonists in the 17th Century – initially referred to as “Hottentots” – were Khoekhoe-speakers. They are usually referred to as the Cape-Khoekhoe and !Ora people (who were previously indicated as Korana [Barnard, 1992]) but their genetic relationships with other extant populations are obscure, as they became “extinct” soon after the arrival of the Europeans. Little is also known about the Taa speaking populations that inhabited the Southernmost area of southern Africa such as the /Xam, the /Xegwi and the Baroa (the latter sometimes referred to as the Mountain bushmen, located in and around the Maloti/Drakensberg mountain range in South Africa/Lesotho), which probably spoke a language similar to the Khomani (of the !Ui group) and who were soon assimilated into Bantu populations who settled in the area. The Karretjie people of South Africa are often considered as the descendants of the /Xam. Given this complex process of contacts and admixture, it is expected that the analysis of admixed populations may help to revive the genetic ancestry of such “vanished” communities and therefore provides a description of the genomic landscape pre-dating the arrival of Bantu speaking populations and European colonists.

Our analysis provides crucial insights into the unsolved histories described above, and more generally on the populations living in the region. Firstly, all of the approaches exploited here point to the existence of an ancient tripartite genetic structure in southern Africa populations, dating back to around 25 Kya (18-32 Kya, Fig. 1B, Fig. 2, Fig. 3, Fig. S5, Fig. S6); these dates are in line with previous estimates for the separation of the two Khoesan components reported earlier [Schlebusch et al., 2012, Pickrell et al., 2012, Kim et al., 2014] and close to the start of Marine Isotope Stage 2 and the beginning of the Last Glacial Maximum, whose impact on the distribution of resources might have triggered such differentiation [Mitchell, 2002]. The first ancestry that we identified –*Northern* Khoesan – mainly comprises Jux’Hoan and !Xun individuals, who live in the Northern Kalahari area. Our TREEMIX and PCA analyses suggest that the two populations are modestly distinct from each other, underlining further structure within this component (Fig. 2B, C). Interestingly, the Khoe-Kwadi speaking Khwe, whose genetic ancestry is mostly Bantu-related, and the Haixxom, share Khoesan genetic affinity with these populations, as expected given their geographic proximity (Fig. 3A). Their genomes also contain the *Central* Khoesan component, which suggests that further admixture with populations with such ancestry may have occurred.

The second ancestry component is common across all the populations in the central Kalahari area (indicated here as the *Central* Khoesan component), and includes all the Taa populations, with the exception of the Khomani (*Southern* Khoesan), but including West and East Khoe-Kwadi speakers and the K’xa speaking population =Hoan, which once again highlights the mismatch between genetics and linguistic affiliation in populations from the region [Schlebusch et al., 2012]. This component has not been reported before, although Schlebusch et al. [2012] reported the unique behavior of Gxui and Gxxana individuals. The inclusion of a more representative set of populations in the current analysis, a few of which are characterized by this Khoesan component, together with a focus on the Khoesan-specific genetic components has led to the secure identification and further characterization of this key element of Khoesan-related ancestry.

The third *Southern* Khoesan component is mainly represented by a set of linguistically heterogeous but geographically proximate populations: the Khomani (Taa speakers), Karretjie, and Nama (Khoe-Kwadi). All of these populations are thought to have originated in the Northern Cape [Barnard, 1992]. Barnard considered “the Khoekhoe and the Bushmen [of the Cape area] as members of a single regional unit, separate from the other (black and white) peoples of the subcontinent” [Barnard, 1992]. This is in agreement with our findings of substantial genetic similarities between these groups, despite their different cultural affiliations. In addition, we found evidence for admixture approximately ~1160 years ago in all the Khoesan populations, suggesting that the arrival of pastoralism happened at the same time across the whole subcontinent.

Taken together, this suggests that cultural diffusion – in the absence of significant gene-flow – might have played an important role in the spread of pastoralism and possibly Khoe languages in southern Africa [Sadr, 1998, Barnard, 2007, Barham and Mitchell, 2008, Schlebusch et al., 2012]. The Khoesan-like genetic ancestry of the Khoe-Kwadi speaking Damara maps to this component, which is consistent with their long-term interaction with the Nama, who speak a very similar language [Guldemann and Fehn, 2014], and possibly coupled with gene-flow from K’xa populations living in the same area (as suggested by the occurrence of the *Northern* Khoesan component in their genetic make-up). All the Coloured and the Bantu populations from the Southernmost part of the continent (South Africa and Lesotho) are characterized by the *Southern* Khoesan component (Fig. 3A), suggesting an overall broad homogeneity in Khoesan ancestry over this specific region. However, it is worth noting that several Bantu individuals in the LAMDS plot are slightly deviated toward central Khoesan populations, and that Bantu populations show substantial differences when compared to Coloured individuals, as our cluster analyses based on MDS and IBS distances suggest. Moreover, consistent differences in admixture times have been detected among the two groups. Given their different geographical distribution, such observations could be explained by the existence of additional Khoesan structure in the region and the past presence of differentiated groups around the Lesotho/Drakensberg area (assimilated by local Bantu speaking groups), or by admixture between Bantu speaking population and groups characterized by *Central* and *Southern* Khoesan ancestries [Busby et al., 2016].

Interestingly, the tripartition observed in the Khoesan ancestry does not recapitulate cultural affiliation (Fig. 3C). As described above, we in fact identified a broad inconsistency between genetic clustering and linguistic or subsistence affiliation [Pickrell et al., 2012]. When we predicted genetic similarity among individuals from geography, predictive error was substantially lower than that of subsistence strategy or linguistic affiliation, both marginally improving the predictive power when considered together with geography. Extensive admixture and cultural transition appears to have characterized populations from this area. Similar scenarios have been proposed also for Europe [Lazaridis et al., 2014, Haak et al., 2015] and Madagascar [Pierron et al., 2014], suggesting a common process across human populations. The importance of geography on the distruibution of the genetic variation among the Khoesan is further confirmed by the geogenetic map inferred by SpaceMix built using random coordinates as prior, which recapitulates the geographic location of the populations.

Our ADMIXTURE analysis of Niger-Congo-speaking populations (which includes Bantu speakers) identified four different ancestral components broadly consistent with their geographic location (Fig. 1B). Specifically, we identified three Bantu components that are represented in Eastern, South-Eastern and Western Africa. Interestingly, the latter is modal in the Damara, and in the pastoralist Bantu-speaking Herero and Himba (from 55% in the Himba to 86% in the DamaraP sample), but not in other Bantu-speaking groups of the region (Mbukushu ~20%, Owambo ~27% and Kwangali ~22%). This component is slightly more related to West Africa than the Eastern and South-Eastern Niger-Congo components, and its differential distribution among Bantu groups in this region may be explained by different waves of Bantu colonists into southern Africa, as suggested in a recent survey of African genetic history based on haplotype analyses [Busby et al., 2016] or possibly related to the Early Iron Age settlers who arrived in the mid-1st millennium AD [Diamond, 1997]. Alternatively, this could reflect the shared demographic history of the Herero, the related Himba and the admixed Damara.

## Conclusions

The genetic structure of southern African populations is complicated by the existence of ancient population structure, onto which several layers of additional genetic ancestries have been overimposed over the last few centuries. Here, we demonstrate that local ancestry approaches can be used to tease apart the genetic structure of such ancient components, characterizing their relationships and current distribution, further supporting a role for widespread admixture in human history [Patterson et al., 2012, Hellenthal et al., 2014, Busby et al., 2015, Montinaro et al., 2015]. Further insights are expected to be collected by the molecular investigation of archaeological human remains [Llorente et al., 2015, Morris et al., 2014]. Beyond the obvious historical and archaeological implications for the reconstruction of the subcontinent dynamics, these observations are of relevance for anthropological studies as well as for epidemiological and translational applications (for example, in the design of genome-wide association studies).

## Methods

### New data

We generated novel genotype data for 59 individuals from seven southern African populations collected in Namibia and Lesotho. Forty-four of these individuals from four Bantu speaking groups (MbukushuM, OvamboM, Kwangali, Sotho) and a Khoesan-speaking group (NamaM) have been published previously [González-Santos et al., 2015], using a subset of markers (~ 2,000). Eight individuals each from the Damara and Haixxom, collected in the Khorixas and Etosha areas of Namibia, respectively, are presented here for the first time. Detailed information about the collecting process and samples are available elsewhere [Marks et al., 2012, 2015, González-Santos et al., 2015]. Full ethical approval for the collections was provided by the Oxford Tropical Research Ethics Committee (OxTREC), the Lesotho Ministry of Health and Social Welfare, the Lesotho Ministry of Local Government, the Lesotho Ministry of Tourism, Environment and Culture, and the Namibian Ministry of Health and Social Services. The Nama, Owambo and Sotho populations were genotyped on the Illumina Human 610-Quad BeadChip (Illumina, San Diego, CA, USA), while the Haixxom, Kwangali, Damara and Mbukushu were genotyped on the Human Omni5-Quad BeadChip (Illumina, San Diego, CA, USA). All the data are available at (https://capelligroup.wordpress.com/data/).

### Existing datasets

Our analyses focus on southern African populations. We therefore merged our data with an additional 31 Khoesan-speaking and 20 “admixed” and Bantu-speaking populations from Li et al. [2008], Consortium [2010], Henn et al. [2012], Schlebusch et al. [2012], Pickrell et al. [2012], Petersen et al. [2013], Pickrell et al. [2014], Lazaridis et al. [2014] (Fig. 1, Fig. S1, Table S1). Additional data from outside of southern Africa were taken from populations with European, African and Middle East ancestry, genotyped on different Illumina platforms and the Affymetrix Axiom Genome-Wide Human Origins 1 array (Fig. 1, Fig. S1, Table S1) [Li et al., 2008, Consortium, 2010, 2012, Patterson et al., 2012, May et al., 2013]. Our final dataset comprised 1,856 individuals from 91 populations (Fig. 1A, Table S1).

### Process for merging datasets

Because the genotype data described above came from multiple different platforms and studies, we performed a systematic pipeline for merging the data, keeping the Illumina and Affymetrix data separated. Each dataset was pre-processed removing markers and individuals with a missing rate higher than 10% using PLINK 1.9 [Chang et al., 2015]. Marker positions were lifted to build 37 human genetic map using data provided by Illumina and Affymetrix and all the non-autosomal markers were excluded from the analysis. Specifically, we first merged all the datasets genotyped on the same platform, discarding individuals and markers with a call rate lower than 98% and excluding SNPs with G/C or A/T mutations, which could lead to errors in the merging procedure. Although, in principle merging genotype data from different platform manufacturer could led to errors or biases, this approach has been successfully employed in previous investigations [Reich et al., 2009, Henn et al., 2012]. In addition, in none of the analysis performed we have seen significant differences between similar or equal groups genotyped with different platforms. Moreover, the IBS similarity between 328 pairs of individuals which have been genotyped by the two manufacturers is always higher than 0.996 (99%*CI* = 0.998 − 1.000). We next used the KING software to infer kinship [Manichaikul et al., 2010], and randomly removed one individual from pairs with a kinship rate higher than 0.0884. The resulting platform-specific datasets comprise 250,547 (Illumina) and 498,140 (Affymetrix) markers, respectively.

### The Unphased Dataset

To maximise the number of populations analyzed (at the expense of SNP density) we merged all data collected from all studies into a large dataset, which we refer to as the “Unphased Dataset”. The two platform-specific datasets, comprising 250,547 (Illumina) and 498,140 (Affymetrix) markers, were merged on physical position to avoid unnecessary loss of markers due to mismatching IDs on different platforms. Following this merge, we again performed the same quality control and removal of relatives described above obtaining a final dataset containing 1,856 individuals genotyped at 63,767 SNPs.

### The Khosean Ancestry dataset

To maintain a high density of SNPs for local ancestry analyses, we analyzed the quality controlled Illumina and Affymetrix datasets separately. For each of the two platform-based datasets described above, we computationally phased the genotype data to generate haplotypes using SHAPEITv2 [Delaneau et al., 2012, 2013] with the human genome build 37 recombination map downloaded from the SHAPEIT website https://mathgen.stats.ox.ac.uk/genetics_software/shapeit/shapeit.html#gmap. We generated a second dataset (“The Khosean Ancestry dataset”), by initially removing from each platform-specific dataset (Illumina and Affymetrix) Non-Khoesan genomic fragments as identified using PCAdmix [Henn et al., 2012]. In brief, PCAdmix builds a PCA space based on reference panels, and projects tested genomic chunks on it, and similar approaches have been previously developed [Omberg et al., 2012, Price et al., 2009, Maples et al., 2013]. Subsequently, the probability of a given ancestry for a selected chromosomal chunk is estimated from PC loadings, and a Hidden Markov Model is then applied to refine them. In the current context, we estimated local ancestry likelihoods in 1 cM windows, using Yoruba, Jux’Hoan, and CEU individuals as ancestry donors. Given the recent West Eurasian genomic component documented in the Jux’Hoan populations as the result of admixture with non-Khoesan populations [Pickrell and Pritchard, 2012, Hellenthal et al., 2014, Busby et al., 2016], only individuals with more than 99% of the “Khoesan component” – as estimated by the *K* = 3 ADMIXTURE run described above – were considered as donors, with the remaining individuals used as target individuals. The final number of Jux’Hoan individuals used as ancestry donors was 28 and 26 in the Illumina and Affymetrix datasets, respectively. To minimise the impact of chunks with mixed ancestry, we post-processed inferred local ancestry estimates by retaining only those windows with a ancestry probability *>* 99%. In addition, we only analyzed individuals characterized genome-wide by more than 35% of the tested ancestry (as for ADMIXTURE analysis for *K* = 3; see below). We tested the accuracy of PCAdmix on the Illumina dataset using a simple approach. In details, using the same source populations (Yoruba, Ju—’hoan and CEU) and parameters, we estimated the Local ancestry of 73 Yoruba individuals. When no threshold confidence was used only the 0.8% of the analyzed 1cM windows were misassigned. However, when only windows assigned with more than 99% confidence has been retained all the misassigned fragments were discarded.

We used a custom-made PYTHON script (*MaskMix*, available at:https://capelligroup.wordpress.com/scripts/), to extract the Khoesan Specific Fragments (KSF) inferred from the post-processing described above.*MaskMix* considers each individual as homozygous and composed by one chromosome per pair only, from which high confidence KSFs were extracted and analysed. This approach allows us to use chromosomal data instead of individual genotypes, maximising the amount of genetic data suitable for the analysis, and is not expected to affect any of the analyses performed because the relative allele frequency would be unchanged. To allow the comparison between individuals genotyped using arrays from different providers, the resulting two datasets were pruned to retain markers which overlapped between the Illumina and Affymetrix datasets and that were located on Khoesan-specific genomic fragments. Finally, we removed all the individuals for which less than 10% of the total number of overlapping SNPs were retained. The resulting dataset is composed of a total of 63,767 markers and 787 individuals. Given the variation in Khoesan ancestry in different individuals, the average number of retained SNPs per individual was 22,442 (median 19,887; range 5,457-50,643). We refer to this final set of SNPs selected as described above as the ”Khoesan Ancestry dataset”. Furthermore, we explored the distribution of the tract length estimated by the approach with respect of the platform manufacturer.

## Statistical Analyses

### Population structure

We applied both model-based and non-parametric clustering approaches to describe population structure in the Unphased Dataset. First, we used the ADMIXTURE [Alexander et al., 2009] maximum likelihood algorithm to estimate the individual-level ancestry, applying the author’s cross validation procedure and a random seed, for all values of *K* = {2…20}. Ten different runs for each value of *K* have been performed, and differents output have been combined using the CLUMMP utility in CLUMPAK [Kopelman et al., 2015, Jakobsson and Rosenberg, 2007] with the LargeKGreedy algorithm, random input order, and 2000 repeats. After post-processing the ADMIXTURE output with DISTRUCT [Rosenberg, 2004], we plotted the results using the *R* statistical programming software and a modified version of *polarHistogram* function from the *phenotypic phorest* package (http://chrisladroue.com/phorest/). For the *K* = 20 run with the highest Likelihood value, we computed pairwise *F*_*ST*_ [Holsinger and Weir, 2009] for each of the *K* ancestral components as implemented by ADMIXTURE, visualizing their distances with a heatmap using the *pheatmap* [Kolde, 2015] R package. Ancestry components were additionally clustered through a complete hierarchical approach [Everitt and Britain, 1980]. For the *K* = 20 analysis with the highest Likelihood ratio, we estimated splitting time between the three components and all the Khoesan populations using the following formula[Holsinger and Weir, 2009, Henn et al., 2012]: 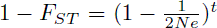 where Ne is the effective populations size and t is the time since separation (in generations). This approach has been applied on admixture ancestries and population pairwise *F*_*ST*_, although in the latter we removed individuals characterized by less than 80% Khoesan ancestry, as reported below. We used the Ne inferred over time for 5 Khoesan by Kim et al 2014, from which we extracted the armonic mean, the mean value and the final population size estimate (10 024, 12 302, 14 024). The density of the splitting times between the three ancestries using the two approaches is shown in (Fig.S10). Principal Components Analysis (PCA) was performed using PLINK 1.9 [Chang et al., 2015]. To focus on the structure of Khoesan populations, we selected only those individuals characterized by more than 80% of the “Khoesan” ancestral component as estimated from the *K* = 3 ADMIXTURE analysis described above; we define the “Khoesan” component as the major ancestry present in Jux’Hoan individuals. We refer to this dataset at the “80% Khoesan dataset”. Populations form the same dataset were tested for admixture using *f*_3_ statistics. [Reich et al., 2010, Patterson et al., 2012] considering all three-populations combinations (3990 combinations, Table S3). We reported all the comparison in Table S3, while significant values are reported in Fig 2B, Fig. S10B. In addition, we performed two different set of *f*_4_analysis, using the qpDstat software and the option ”f4mode=YES’. In details we performed the *f*_4_ stat in the form *f*_4_(*Ju|′hoanP e, Nama, X, Chimp*) where X is represented by all the other populations in the dataset (Fig.S10A). Moreover, we assessed the f4 test of the form *f*_4_(*Khoesan*1*, Khoesan*2*, X, Chimp*), where X is one of the admixed (Basters and Coloured) or Southeast Bantu population (Fig.S10C).

### Evaluating ADMIXTURE performance on simulated samples

As we describe in the results, at *K* = 14, ADMIXTURE identify an *European* ancestral component which characterise the admixed populations (Coloured and Basters). However, the *F*_*ST*_ distance between this component and the African ones (Fig.5 is smaller than the distanc between the African and the other European ancestries, which could suggest that the algorithm picked a new generation of allele frequencies caused by the admixture. To test this hypothesis we evaluate the performance on admixture on simulated admixed samples. In details, we used the Yoruba (YRI) and British (GBR) genetic data in Busby et al. [2016] and split the samples in two equal size groups, *”Source”* and *”Admixture”*. The first has been used to generate 4 groups of 25 admixed individuals composed by a variable fraction (20% −40% −60% −70%) of British and Yoruba individuals, which admixed N generations ago with *N* = 5, 10, 30, 50, 70, 100. The admixed individuals have been combined with the *”Admixed”* group. The resulting datasets have been used to perform an ADMIXTURE run for *K* = 2 and *K* = 3 (Fig.S6).

### TREEMIX Analysis

A maximum likelihood tree describing the relationships between Khoesan populations was inferred using allele frequency distributions implemented in the TREEMIX software [Pickrell and Pritchard, 2012, Pickrell et al., 2012]. Given the high complexity of the original dataset, we selected 35 populations to represent all the Khoesan populations and a subset of African and European populations (“TreeMix populations”). We used a chimpanzee outgroup using genome data available in [Patterson et al., 2012], and accounted for LD by jack-knifing over blocks of 500 SNPs, as suggested by the authors in [Pickrell and Pritchard, 2012]. The robustness of the resulting tree was tested by performing 100 bootstrap runs and estimating branch support using DENDROPY software [Sukumaran and Holder, 2010].

We performed ten different runs using different random seeds (Fig.S9), and report the tree with the maximum support in Fig.2B. To visualize only the Khoesan ancestry and to remove the confounding factors of admixture, we informed TREEMIX of existing relationships between Khoesan and non Khoesan populations using the cor mig and climb commands (Table S2). The amount of genetic ancestry shared between populations was estimated through the *K* = 3 ADMIXTURE run described above. It is important to note that these estimates are not fixed values, but are used by the algorithm as starting points to infer the maximum likelihood estimates [Pickrell et al., 2012].

### Population structure inference using the Khosean Ancestry dataset

Referring to the Khosean Ancestry dataset, we estimated pairwise (1-IBS) genetic distance with PLINK v1.9, correcting for missing data, and summarized relationships with a Multi-Dimensional Scaling (MDS) plot, using the *cmdscale* function in *R* (Fig. 2A, Fig. 3A). We corrected the inferred IBS-based distances using the formula 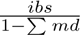, where *md* is the number of missing data in the pair of individuals analyzed. Furthermore, after visual inspection, we removed 25 outlier chromosomes (1 chromosome each from AmaXhosa, Basters, ColouredEC, ColouredWellington, Khwe; 2 from SEBantu and Sotho; 3 from Tswana and Kgalagadi, 8 from SouthAfricaBantuMa). Distances are computed using the number of SNPs shared across pairs of individuals, which differs across pairs given the variation across individuals in the number of SNP markers found on khoesan genomic fragments. The average number of markers used for individual to individual comparisons is 9,310 (median= 6404, range= 111-46,419). In order to assess the bias that a small number of SNPs may cause in capturing the genetic variation in the area, we resampled 10 different datasets composed by N markers, with *N* = {500‥1000…10000} and compared the median and 95% confidence interval with the whole dataset. In addition, reported the average *R*^2^ between the resampled and the full datasets.

### Structure and distribution of Khoesan ancestry in southern African populations

To assess the presence of ancient structure in Khoesan speaking populations and the existence of different Khoesan ancestry in admixed Bantu and Coloured individuals, we explored the MDS coordinates and IBS distance matrix. We initially plotted all southern African groups with 90% utilization distribution density kernels for the three ancestries, estimated using the Kernel Utilization Distribution (UD) function in the *adehabitatHR* R package [Calenge, 2006]. The function estimates the minimum area of the plot in which individuals from the same population have the 90% probability of being located. Firstly, we used the MDS coordinates and IBS distances to group all individuals into *N* different clusters, for values of *N* = {1…9}, using the algorithm implemented in Mclust R package [Fraley and Raftery, 2002]. We next visualized the average assignment probabilities into population- and affiliation-based barplots. Finally, to test for a correlation between genetic and geographic distances, we performed a Procrustes test [Peres-Neto and Jackson, 2001] as implemented in the R package *ade4* [Dray and Dufour, 2007, Dray et al., 2007], with 1000 bootstrap iterations between geographic and MDS (genetic) coordinates.

### SpaceMix analysis

In order to further investigate the distribution of the genetic variability among the three ancestries previously described, we built a ”GeoGenetic map” of Khoesan populations (only individuals with more than 80% Khoesan ancestry), taking advantage of the Bayesian statistical framework implemented in SpaceMix [Bradburd et al., 2016]. Briefly, this approach reconstructs the genetic relatedness among populations as a map in which distances are proportional to their genetic dissimilarities. Moreover, inferred long distances relatedness are modelled as gene flow between populations. In details, we have run an analysis as in Bradburd et al. [2016]: first, five independent short chains of 5 * 10^6^ Markov Chain Monte Carlo interactions, in which only locations were estimated were run. For the whole analysis, the initial population locations were taken by a uniform distribution with minimum and maximum of 180,-180 and 90,-90 for longitude and latitude, respectively. Secondly, a long chain of 10^8^ iterations sampled every 10^5^ steps has been analyzed. The starting parameters of this chain were taken by the last iteration of the short run characterized by the highest posterior probability. Finally, the inferred ”geogenetic position” (and their 95% Confidence interval ellipses) and and their sources of admixture were superimposed on observed population sampling locations (Fig. 13A). The overall performance of the approach has been assessed exploring the posterior probability trace, (Fig. 13B), while the ability of the model to describe the data has been evaluated analyzing the correlation between parametric vs. observed co-variance matrix (Fig. 13C, D) and the decay of co-variance versus geographic distance for observed and inferred matrices.

### Estimating predicting power for geographic location, linguistic affiliation and type of subsistence

To assess the role of geography, subsistence and language in predicting genetic variation of Khoesan populations, we performed regression analysis of LAMDS first two components versus all the other variables, singularly or combined (”Geography + Language”, ”Geography + Subsistence” and ”Geography + Language + Subsistence”; Table S4). All the combinations were tested trough a 5-folded cross validation analysis in which the dataset was split in 5 random subset. Each of these subsets was then tested against the other four and the combined error recorded and showed in a barplot (Fig. 3C). Two alternative subsistence affiliation lists were used to take into account uncertainty in designation or the co-existence of multiple subsistence strategies. In details ”Subsistence 1” was annotated according to Schlebusch et al. [2012] and Barnard [1992]. In order to take into account the multiple and/or uncertain subsistences in Damara and admixed populations, we used an alternative list (”Subsistence 2”) in which these groups were indicated as Hunter-Gatherer/Herder and farmers, respectively (Table S4).

### Estimating admixture dates using MALDER

We assessed possible evidence for admixture using the algorithm implemented in MALDER [Pickrell et al., 2014], which, developed from ALDER, fits a mixture of exponential functions to weighted Linkage Disequilibrium density curves, allowing to identify multiple admixture events. We performed a MALDER analysis for all the populations with more than 2 individuals using the *mindis* : 0.005 parameter. The results of our analysis are shown in Figure S9, in which we report the estimated dates (±1*SD*) and the two populations generating the highest amplitude for each inferred event. In addition, for each event we assessed if the other amplitude estimates were significantly different than the maximum one (*Z* > 2). Z has been estimated using the formula:

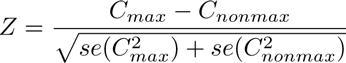

We classified the two populations which generate non significant *Z*_*score*_ (*Z* < 2) into one of the macrogroups showed in Fig 1A and represented as a barplot showing the frequency of comparisons which show that group. Each dataset from different platform manufacturer has been analyzed singularly. Although the overall interpretation of the results can still be challenging, this could be improved by this approach.

## Acknowledgments

This work was funded by a Leverhulme Trust Research Project Grant (”The genetic landscape of southern Africa human populations”) and supported by the Wenner-Gren Foundation, the University of Oxford Boise Fund, the John Fell Oxford University Press (OUP) Research Fund, and Fundação para a Ciência e Tecnologia (grant number SFRH/BD/90648/2012 to M.G.-S.). We thank all the people who donated their DNA samples making this work possible and all the various people and institutions that helped with the organization of the fieldwork and the collection of the samples. We are grateful to Joe Pickrell and David Reich for the sharing of southern Africa genetic data; Hie Lie Kim and Stephan C. Schuster for making available their PMSC results. We are grateful to Peter Mitchell for the suggestions on archaeological patterns in southern Africa.

## Authors contributions

F.M. and C.C. conceived the study; F.M., G.B.J.B and M.G.S. performed the analyses; O.O. and E.O. provided support for the collection of DNA samples; P.A., G.D.B. and V.P. contributed to the genotyping of a subset of the novel samples presented here; F.M, G.B.J.B and C.C. wrote the manuscript with the contribution of all the other authors. All the authors read and approved the manuscript.

## Tables

Table S1 – Populations used in this study. Information on number of individuals in the ”Unphased dataset”, TREEMIX dataset, and dataset with 80% Khoesan ancestry, number of chromosomes in the Khoesan Ancestry dataset, geographic coordinates, group and data source is reported.

Table S2 – Migration events incorporated in the TREEMIX analysis. The “Recipient” population is expected to have a percentage (“Amount”) of ancestry from the “Source” population.

Table S3 – Three-population test (f3 test) results between “Target” and sources (“P1” and “P2”) populations, including standard deviation (sd) and Z-score.

Table S4 – Geographic, Linguistic and subsistence affiliation used in the correlation analysis of MDS coordinates.

**Supplementary Figure 1:**
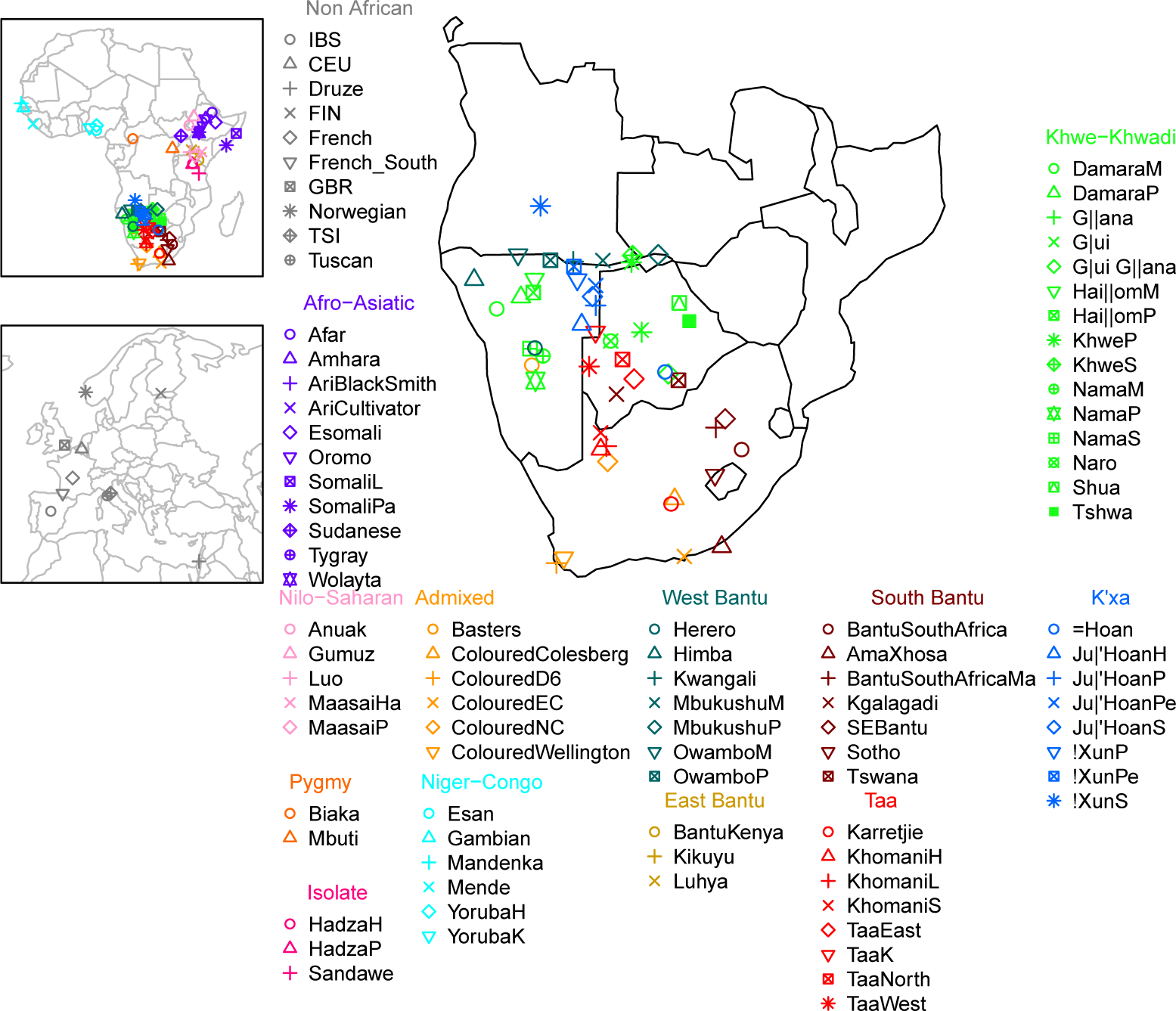
Geographic locations of analysed populations. We analyzed 1856 individuals from 91 groups, genotyped with different Illumina platforms and the Affymetrix Axiom Genome-Wide Human Origins 1 array.

**Supplementary Figure 2:**
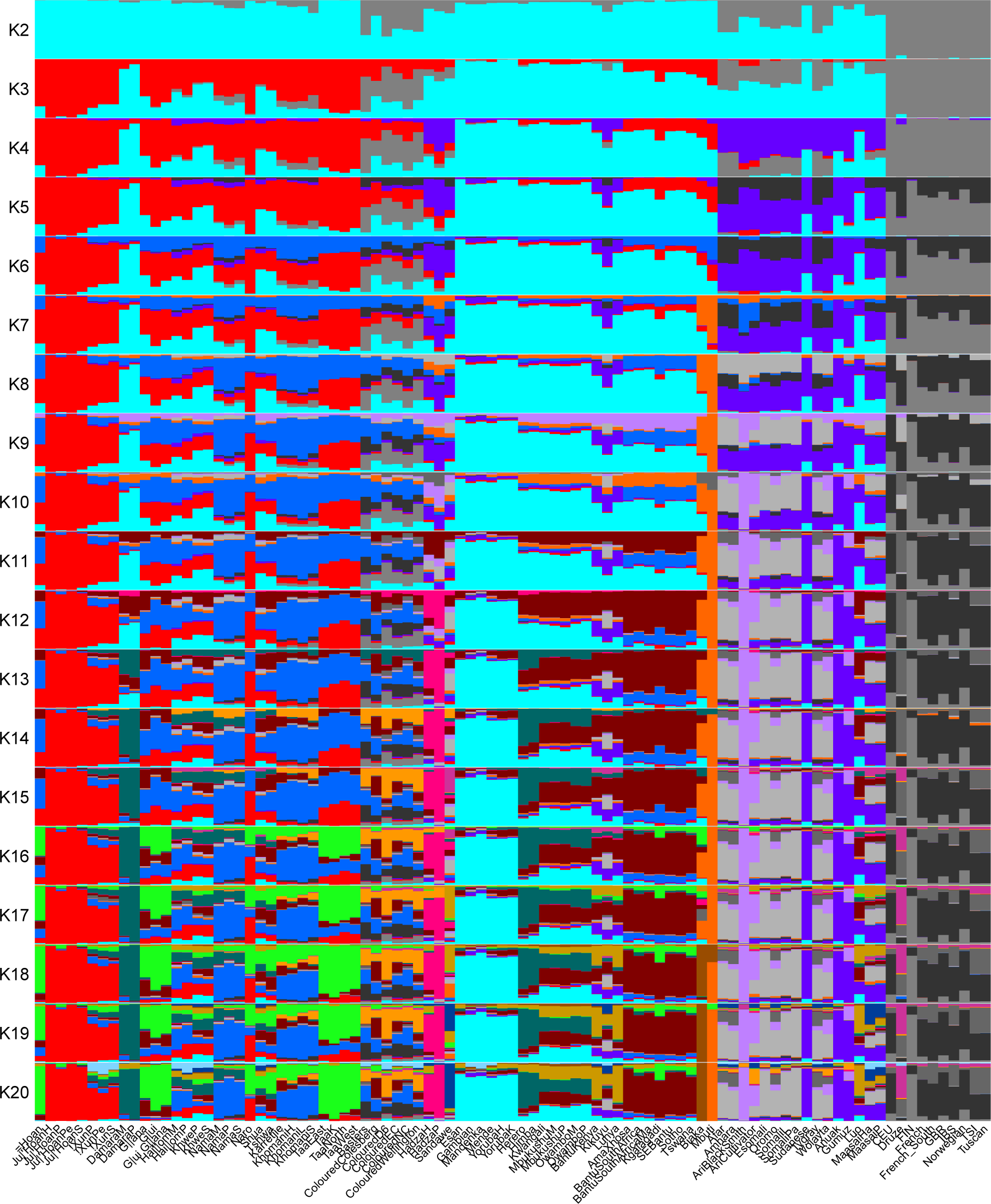
ADMIXTURE analysis of analysed populations for *K* = 2..20. We performed ADMIXTURE analysis, with different random seeds for all the individuals and reported the population average.

**Supplementary Figure 3:**
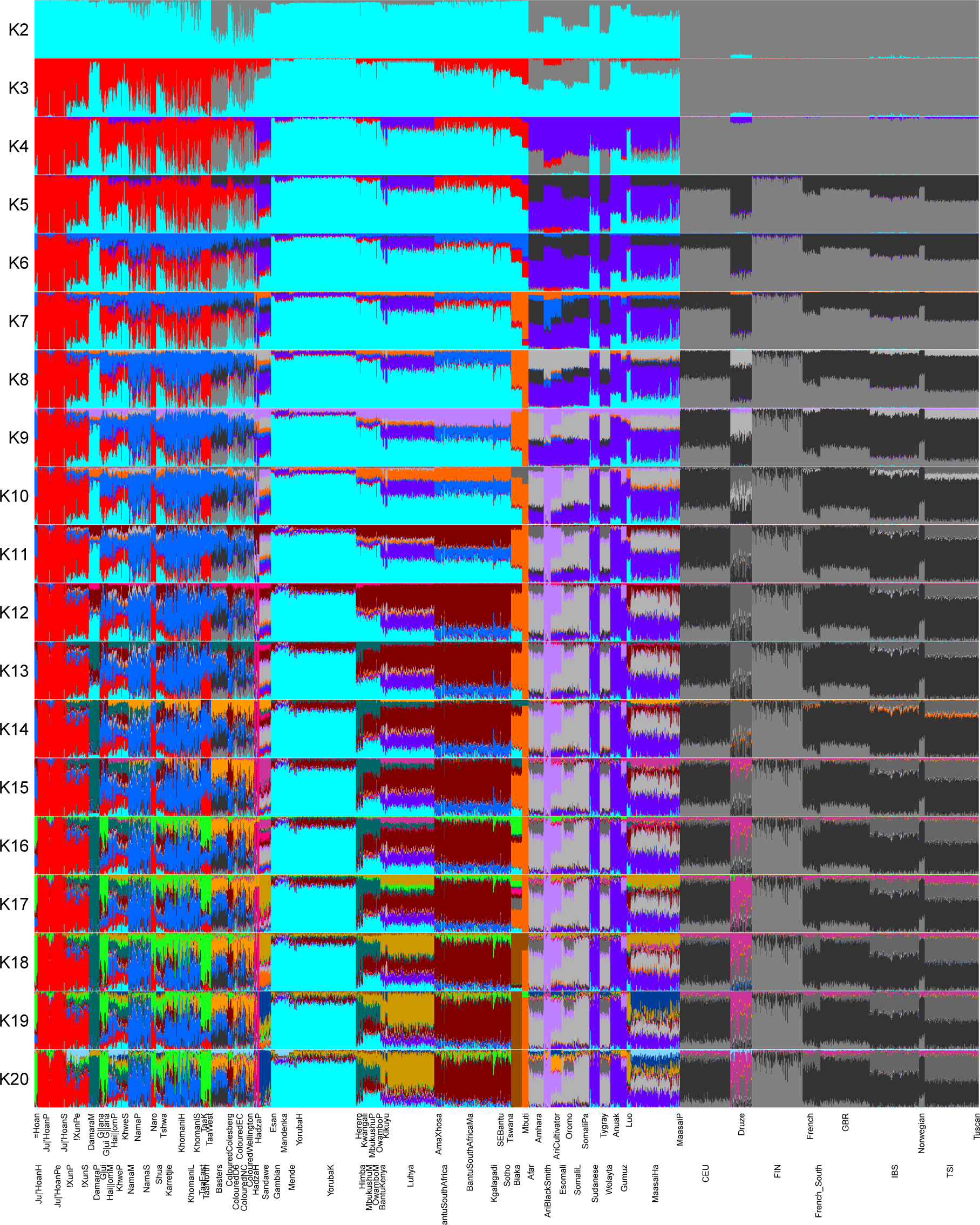
ADMIXTURE analysis of analysed populations for *K* = 2..20. We performed ADMIXTURE analysis, with different random seeds for all the individuals and reported the individual clustering assignation.

**Supplementary Figure 4:**
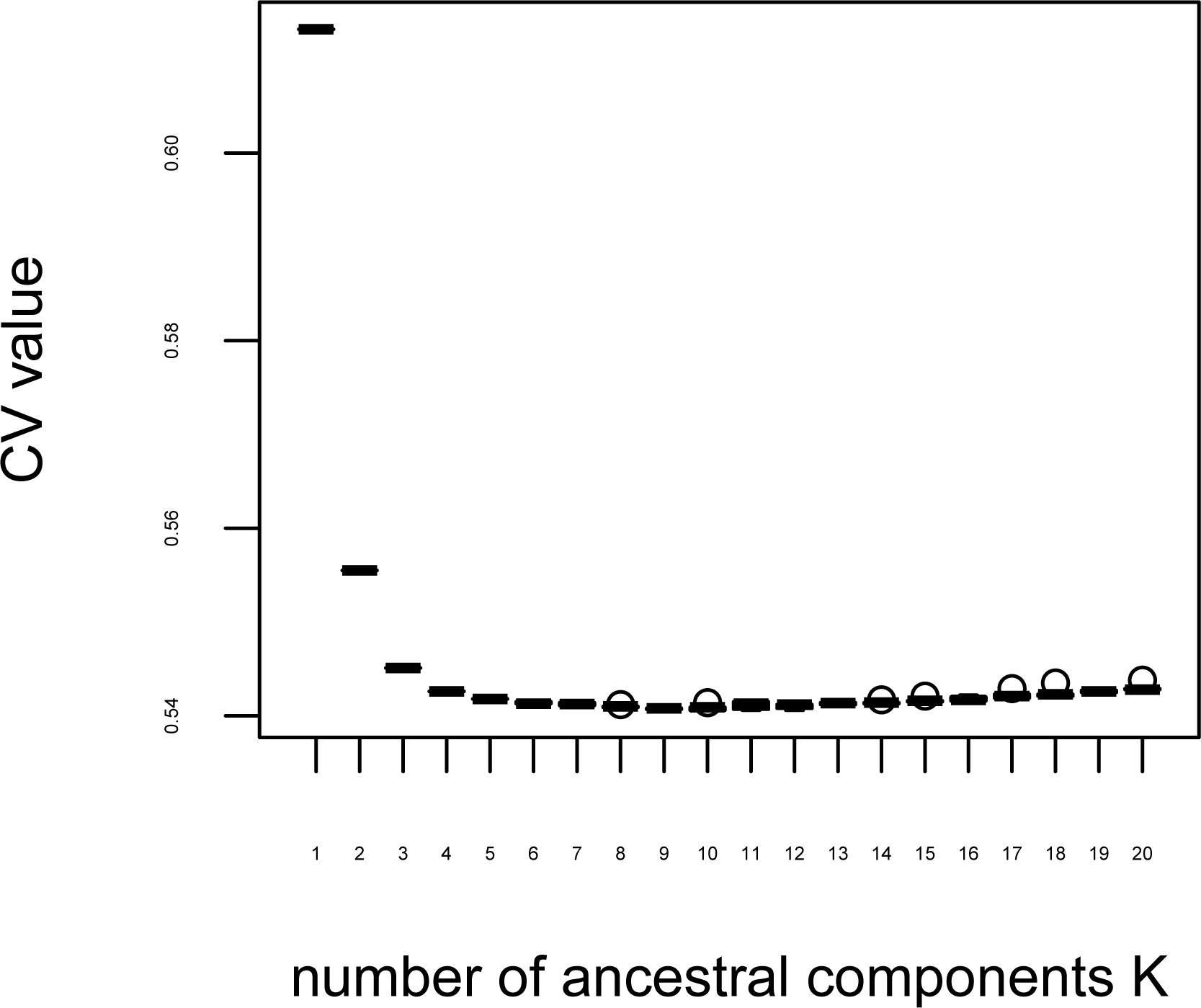
Cross validation error values for ADMIXTURE run for K=2..20. The lowest values for CV error are between 9 and 10 clusters. We noted that analysis for larger number of K provided important insights into the genetic structure of analyzed populations and discussed these in the main text.

**Supplementary Figure 5:**
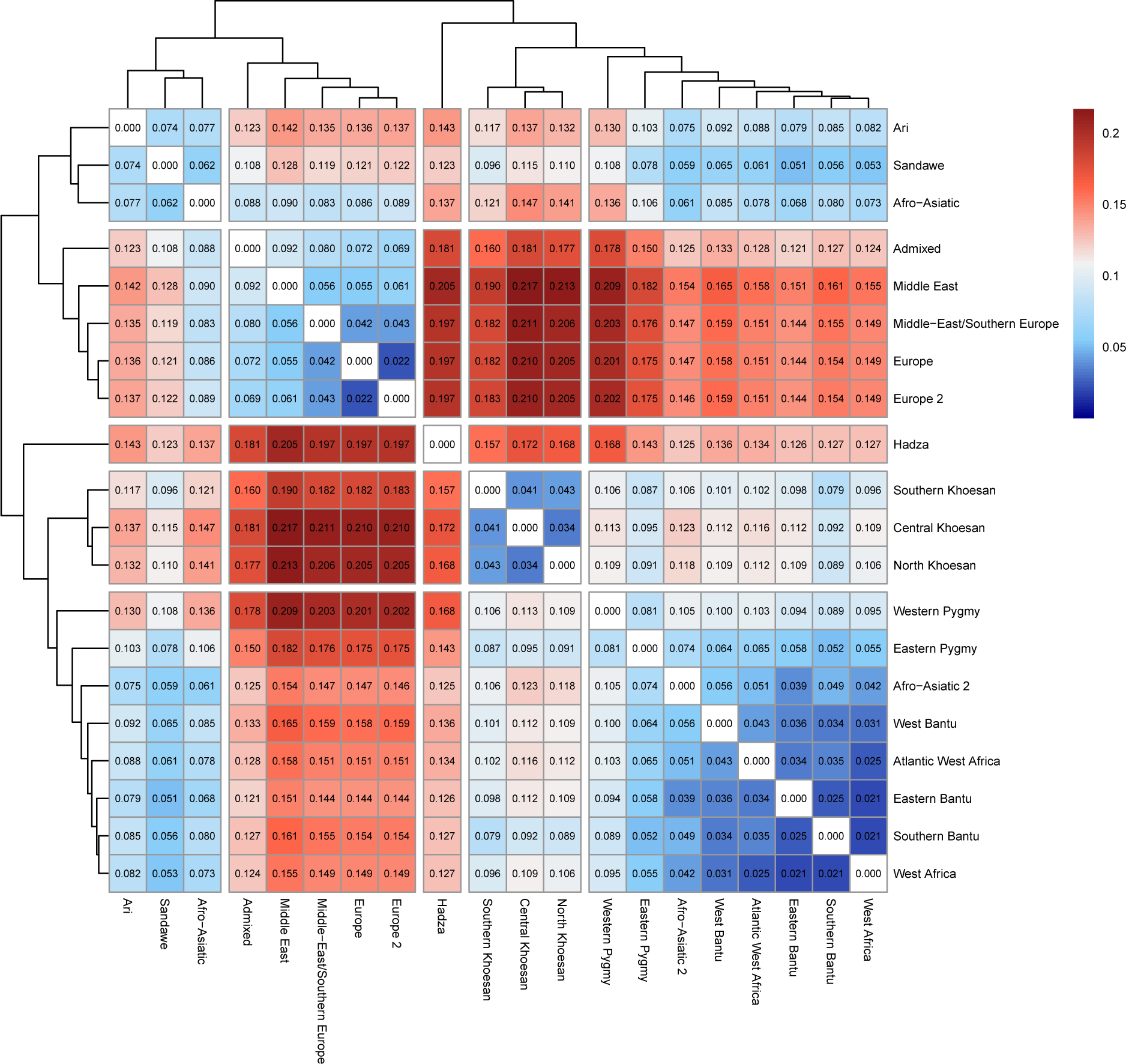
Heatmap and hierarchical trees of the genetic relationships between ancestral components. We reported *F*_*ST*_ values between components (*K* = 20), and clustered them using a hierarchical approach as indicated in Methods section.

**Supplementary Figure 6:**
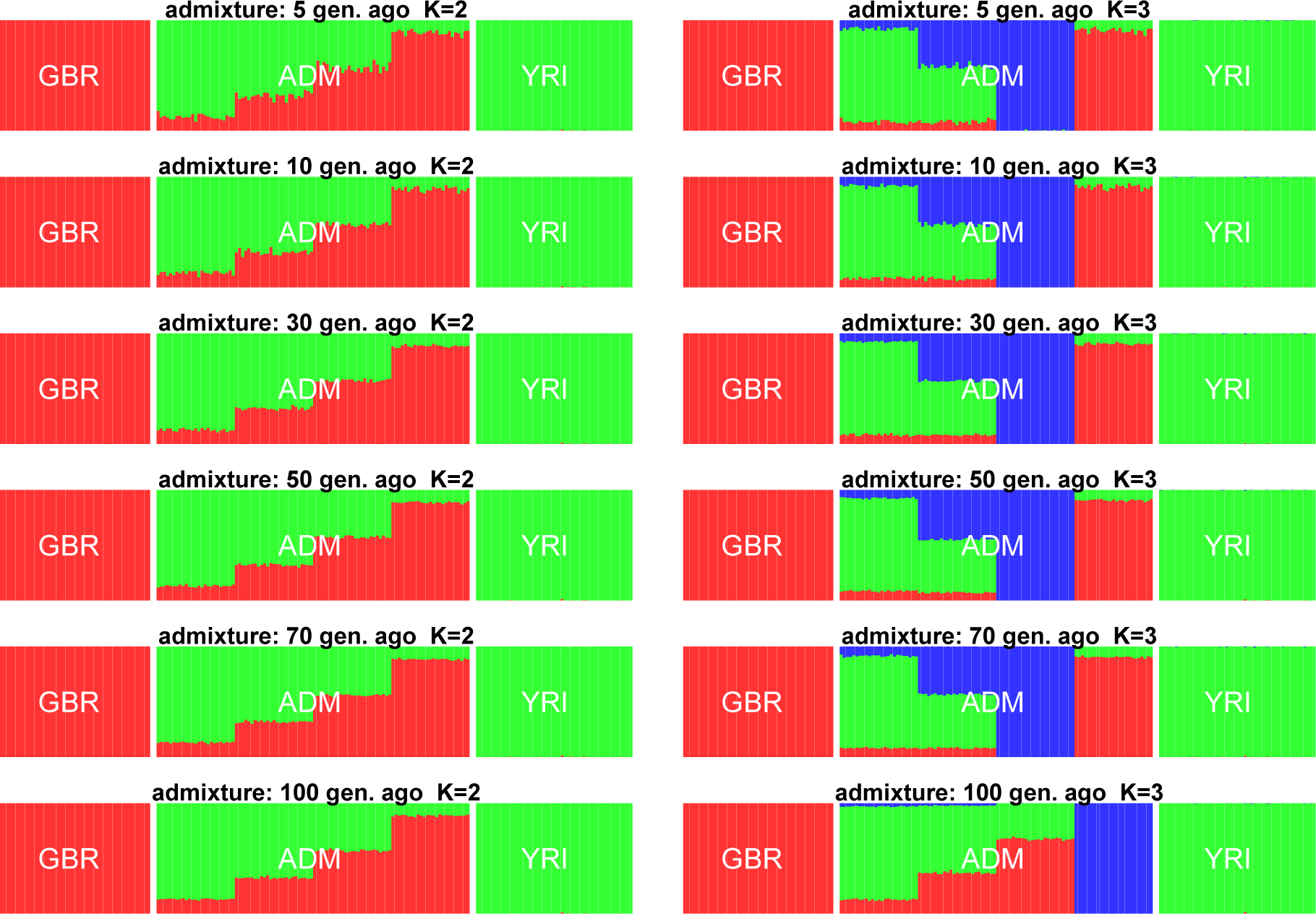
ADMIXTURE analysis of simulated admixed populations for *K* = 2 − 3. We have simulated admixed individuals (ADM) composed by a variable proportion (20% − 40% − 60% − 80%) of British (GBR) and Yoruba (YRI) ancestry. We simulated recent (5 − 10 − 30 generations ago) and ancient (50 − 70 − 10 generations ago) dates of admixture and performed an ADMIXTURE run for *K* = 2 and *K* = 3.

**Supplementary Figure 7:**
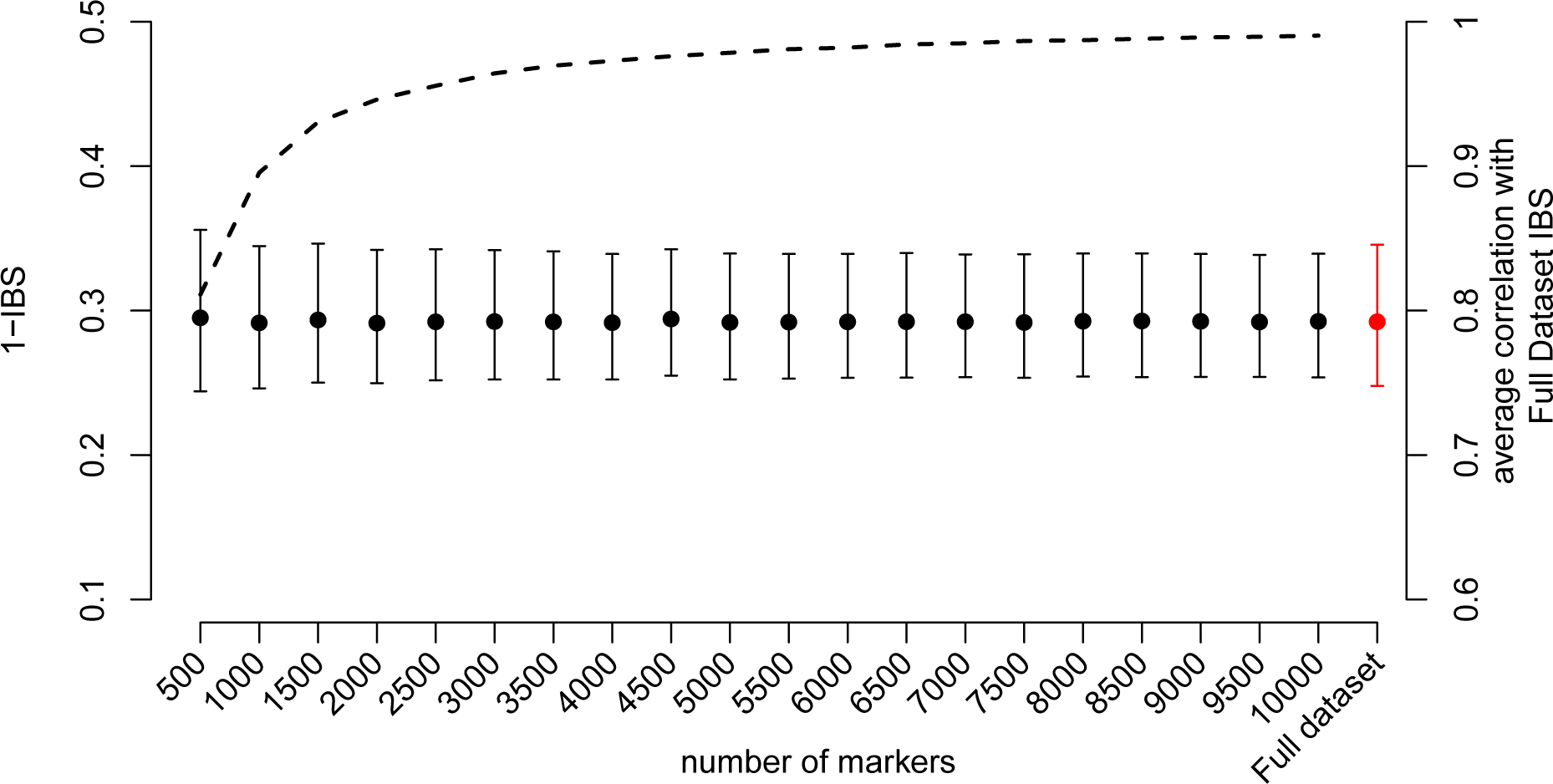
Evaluation of the impact of the number of markers on IBS distance estimation: We resampled 10 datasets composed by N markers, with *N* = {500…1000…10000} and reported the median and the 95% CI values. In addition, for each N the average *R*^2^ coefficient of determination has been reported (dotted line, right vertical axes)

**Supplementary Figure 8:**
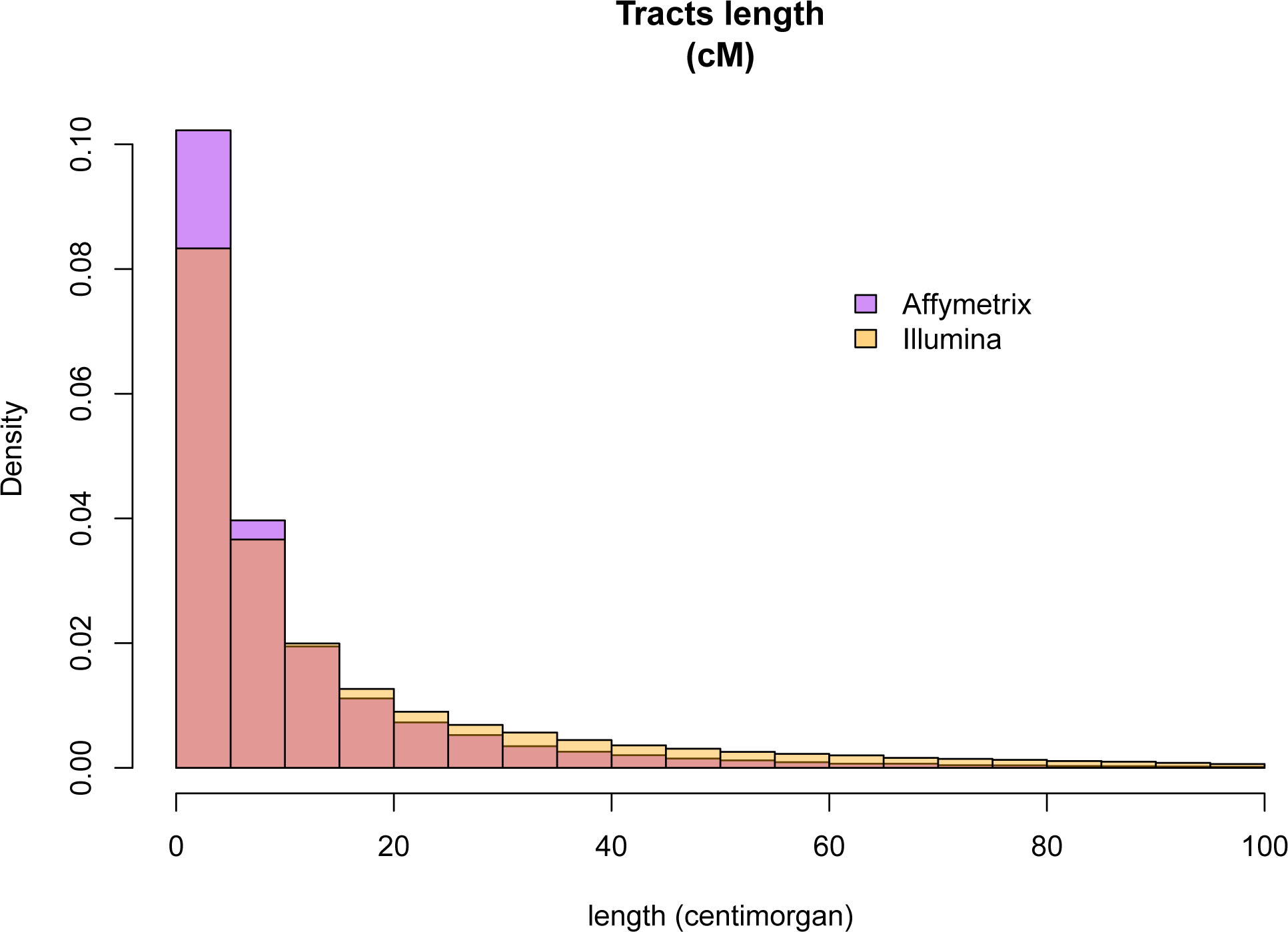
Distribution of the Khoesan tracts length identified by PCAdmix. We estimated the length in centimorgan for the Illumina and Affymetrix dataset. The two distribution are very similar although a higher number of short tracts have been identified for the Affymetrix platform

**Supplementary Figure 9:**
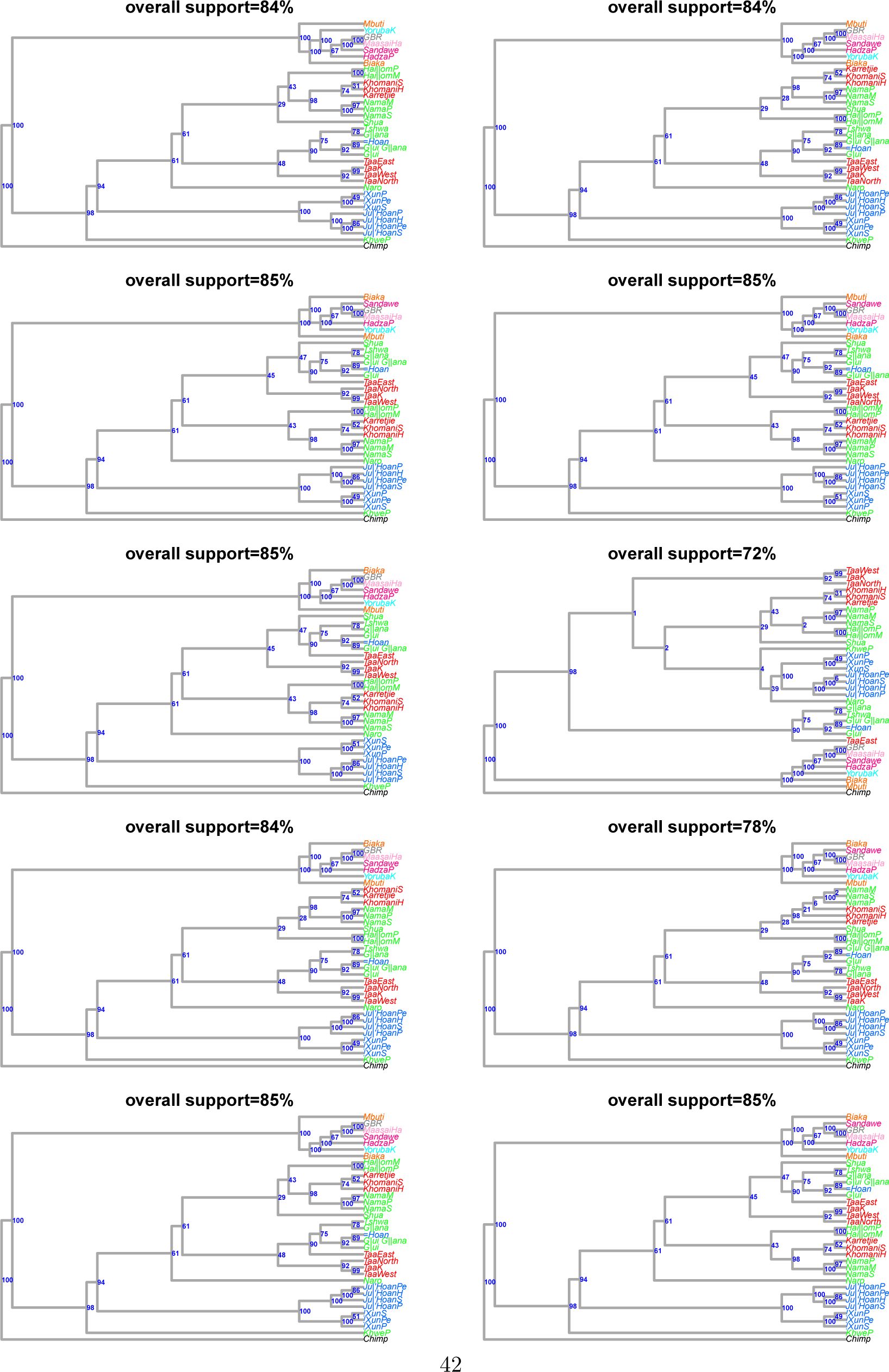
Maximum Likelihood trees of Khoesan populations. We performed ten different runs, using different random seeds of TREEMIX, and assessed their support trough 100 bootstrap replications. We then reported one of the most supported trees in Fig 2B. Note that the most supported trees are identical in their topology.

**Supplementary Figure 10:**
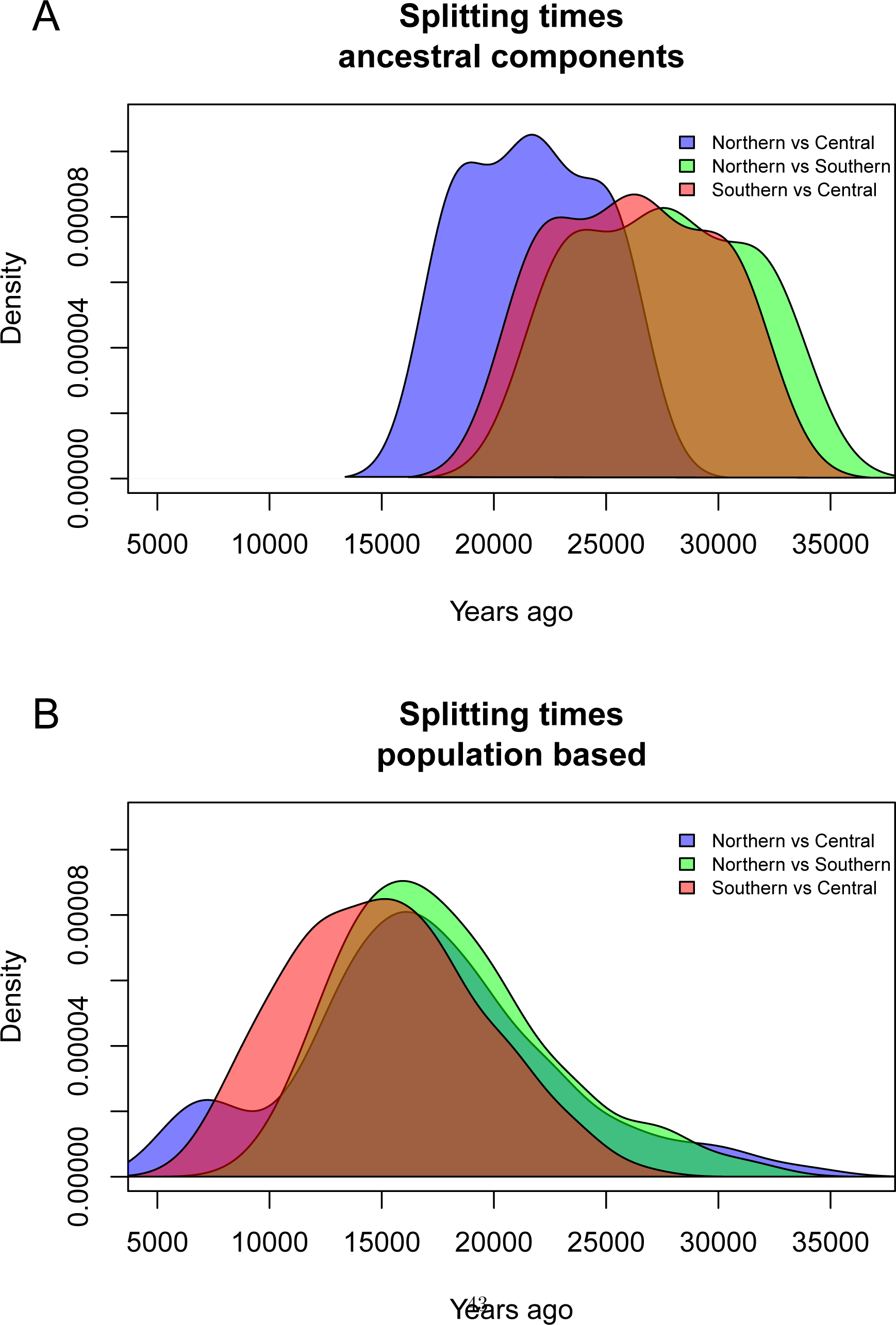
Estimates of the time of the split between the three Khoesan groups. We estimated the pairwise *F*_*ST*_ value between ancestral components (A) or populations (B) and converted each value in *splitting time* using the formula 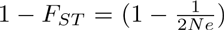 where Ne is the effective populations size from [Kim et al., 2014] and t is the time since separation (in generations).

**Supplementary Figure 11:**
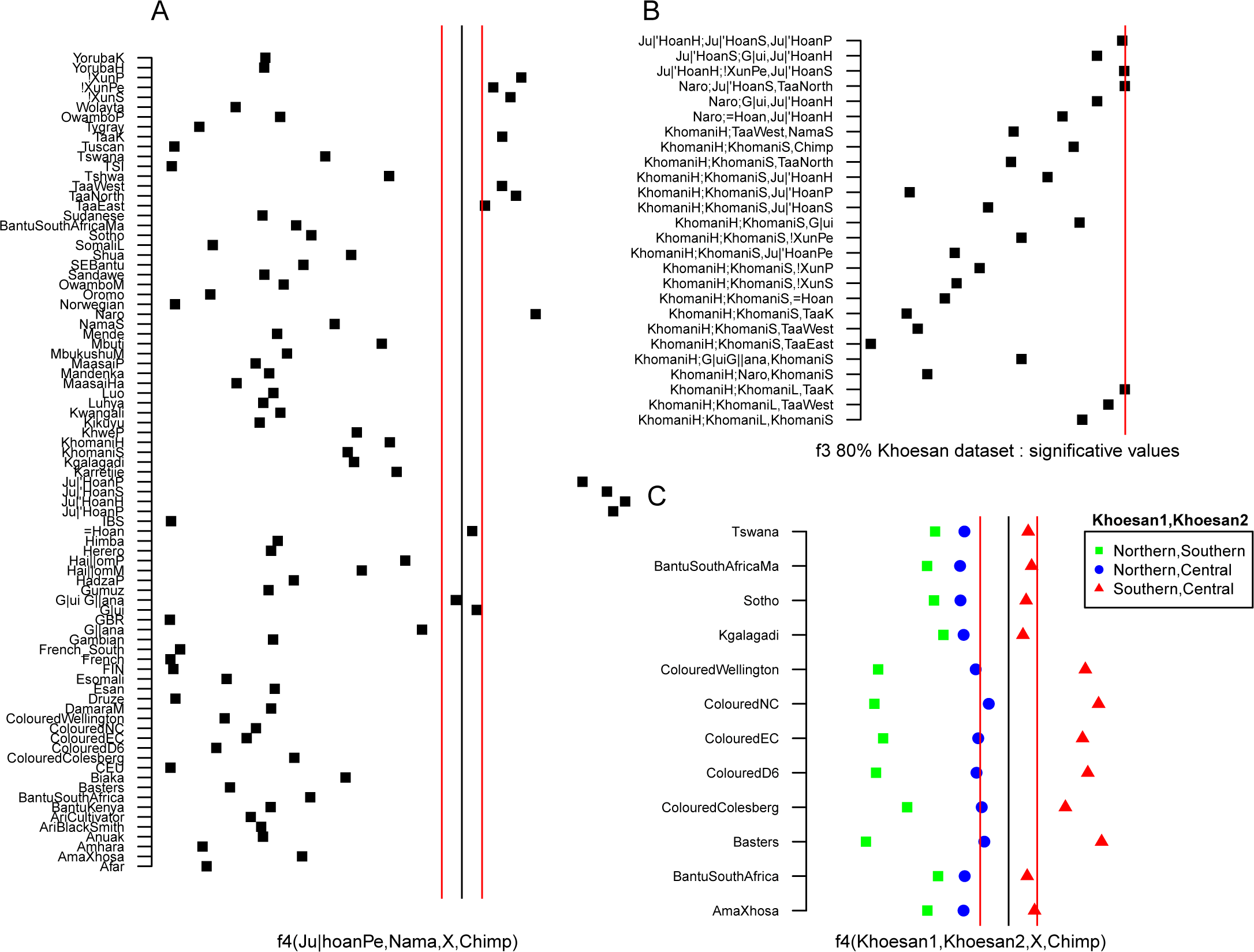
*f*_3_ and *f*_4_ tests A. We first performed a test on the form *f*_4_ (*Ju*|′*hoan, Nama*, *X*, *Chimp*) in order to verify if the *Central* Khoesan group is a mix between the other two groups. Some but no all of the *Central* populations are closer to the Jul’hoan. B. Second, we perform a *f*_3_ test among all the triplettes on the Khoesan populations (including only individuals with more than 80% of Khoesan ancestry. With the exception of the Naro population, none of the *Central* groups show significant (*Z* < −3) values. C. In order to obtain further insights into the Khoesan legacy in Coloured and Bantu populations we performe a *f*4(*Khoesanl*, *Khoesan2*, *X*, *Chimp*) in which Khoesan 1 and Khoesan 2 was represnted by Jul’hoan, Glui & Gllana, and Nama. The Coloured seem to be more similar to the *Southern* populations, while the Bantu seem to be equally similar to the *Central* and the *Southern* (although the Z score is positive, denoting a higher affinity with the *Southern*)

**Supplementary Figure 12:**
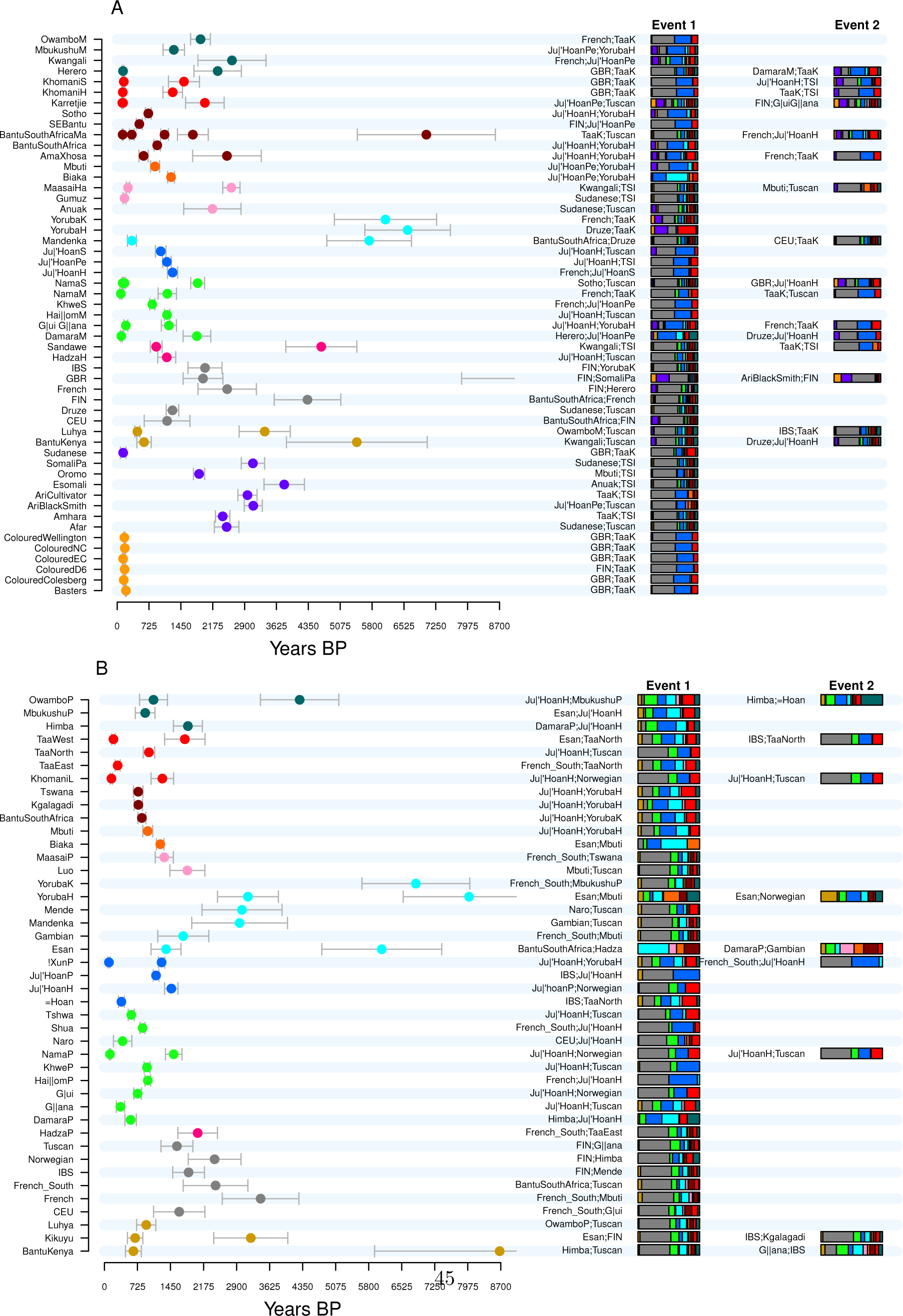
Date of admixture estimates using MALDER. For each platform (A. Illumina and B. Affymetrix) we estimated the date of admixture using MALDER fitting a mixture of exponential functions to weighted Linkage Disequilibrium density curves, estimating multiple admixture, and reporting the best fit. In order to provide a better interpretation of the putative admixture sources, we shows the fraction of populations from a certain area which are non significantly different to the best fit.

**Supplementary Figure 13:**
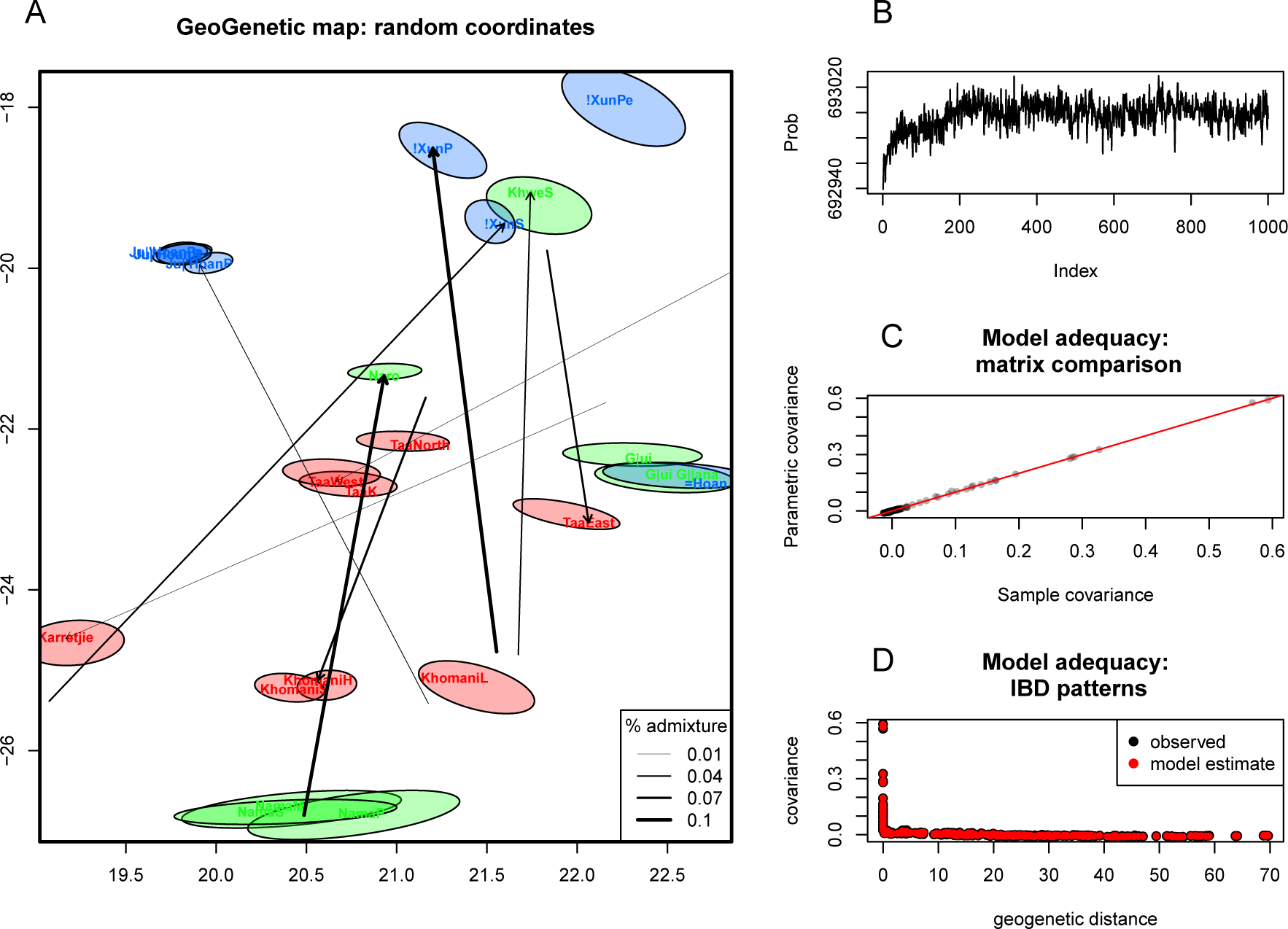
SpaceMix analysis confirms the importance on geopgraphy into the genetic structure of Khoesan populations A. We built a GeoGenetic map of the Khoesan populations using SpaceMix. Firstly, We have run an analysis composed by five independent short chains of 5 * 10^6^ Markov Chain Monte Carlo interactions, in which only locations were estimated. For the whole analysis, the initial population locations were taken by a uniform distribution with minimum and maximum of 180,-180 and 90,-90 for longitude and latitude, respectively. Secondly, a long chain of 10^8^ iterations sampled every 10^5^ steps has been analyzed and the 95% Confidence interval ellipses are shown. B. The performance of the approach has been assessed exploring the posterior probability trace while the ability of the model to describe the data has been evaluated analyzing the correlation between parametric vs. observed co-variance matrix (C.) and (D.) the decay of co-variance versus geographic distance for observed and inferred matrices.

**Supplementary Figure 14:**
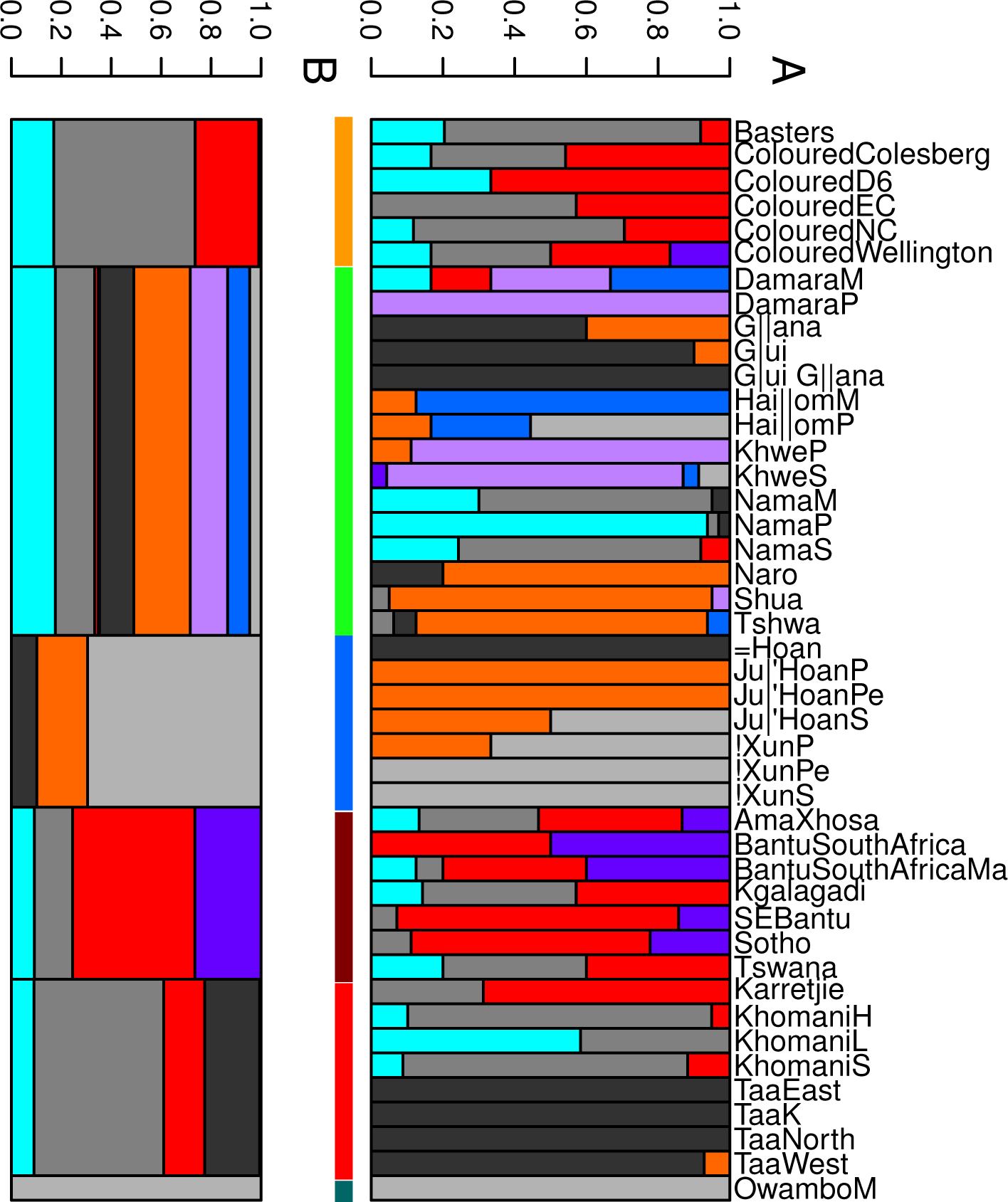
Local ancestry MDS clustering analysis reveals Khoesan-related structure in admixed populations Cluster Analysis of Multi-Dimensional Scaling coordinates. We grouped all the individuals in 9 clusters, using the ibs distance matrix between individuals, as inferred by mclust R package (see methods) and visualized the results in barplot according to populations and language/ethnic affiliation. Colour keys are as in Fig.1A and Fig S1. The results highlight the large heterogeneity in populations from the same affiliation and the existence of a slight but significant substructure between Bantu and Coloured populations.

